# PP2A-B55^SUR-6^ promotes nuclear envelope breakdown in *C. elegans* embryos

**DOI:** 10.1101/2023.06.11.544482

**Authors:** Sukriti Kapoor, Kuheli Adhikary, Sachin Kotak

## Abstract

Nuclear envelope (NE) disassembly during mitosis is critical to ensure faithful segregation of the genetic material. NE disassembly is a phosphorylation-dependent process wherein mitotic kinases hyper-phosphorylate lamina and nucleoporins to initiate nuclear envelope breakdown (NEBD). In this study, we uncover an unexpected role of the PP2A phosphatase B55^SUR-6^ in NEBD during the first embryonic division of *Caenorhabditis elegans* embryo. B55^SUR-6^ depletion delays NE permeabilization and stabilizes lamina and nucleoporins. As a result, the merging of parental genomes and chromosome segregation is impaired. This NEBD defect upon B55^SUR-6^ depletion is not due to delayed mitotic onset or mislocalization of mitotic kinases. Importantly, we demonstrate that microtubule-dependent mechanical forces synergize with B55^SUR-6^ for efficient NE disassembly. Finally, our data suggest that the lamin LMN-1 is likely a bona fide target of PP2A-B55^SUR-6^. These findings establish a model highlighting biochemical cross-talk between kinases, PP2A-B55^SUR-6^ phosphatase, and microtubule-generated mechanical forces in timely NE dissolution.

## Introduction

Mitotic entry and its progression is a complex process, and a significant change in the phosphoproteome is required to orchestrate these events. Multiple kinases act together to bring about these changes and ensure that the duplicated genome is equally partitioned to the daughter cells post-mitosis. In metazoans, one pivotal step during mitosis onset is the nuclear envelope breakdown (NEBD), which allows microtubules to capture sister chromatids and assemble the mitotic spindle (reviewed in Cohen-fix and Askjaer, 2017; Pintard and Bowerman, 2019; Ungricht and Kutay, 2017; Kutay et al., 2021). NEBD involves the systematic disassembly of the gigantic nuclear pore complexes (NPCs) of around ∼100 MDa and depolymerization of the lamina leading to the retraction of the nuclear envelope into the mitotic endoplasmic reticulum (reviewed in Ungricht and Kutay, 2017; Kutay et al., 2021). A tremendous effort in the past decades has linked several mitotic kinases, including cyclin-dependent kinase 1 (CDK-1), polo-like kinase 1 (PLK-1), Aurora A kinase, and NIMA-related kinase (NEK) to NEBD (Heald and McKeon, 1990; Peter et al., 1990; Portier et al., 2007; Hachet et al., 2007; Laurell et al., 2011; Hachet et al., 2012; Linder et al., 2017 Rahman et al., 2015; Solc et al., 2015; Linder et al., 2017; Martino et al., 2017; Velez-Aguilera et al., 2020; Nkoula et al., 2023). These kinases promote hyperphosphorylation of the lamina and nucleoporins (NPPs), which are the subunits of NPC, resulting in loss of nuclear integrity and mixing of nuclear and cytosolic contents during mitosis.

In addition to biochemical changes brought about by these kinases, centrosomal microtubule and the nuclear envelope (NE) localized dynein-dependent mechanical forces are implicated in generating tension that helps in NEBD (Beaudouin et al., 2002; Salina et al., 2002). However, whether kinases are the sole drivers of biochemical changes that help NEBD or this process also relies on additional modulators remains to be discovered. Also, if microtubule and dynein-dependent forces play a role in initiating NEBD beyond mammalian cells remains unknown.

Other than kinases, mitotic progression is equally dependent on the function of phosphatases. It is now viewed that the dephosphorylation of substrates by the phosphatases is as tightly controlled as the phosphorylation reaction by the kinases, and thus, the balance action between kinases and phosphatases sets the level of phosphorylation on various substrates, which is critical for mitotic progression (reviewed in Barr et al., 2011; Gelens et al., 2018; Moura and Conde, 2019; Nilsson, 2019). PP2A is one of the major phosphatases involved in myriad aspects of mitosis, for instance, kinetochore-microtubule interaction, spindle orientation, centrosome disassembly, and cytokinesis (Foley et al., 2011; Kruse et al., 34; Suijkerbuijk et al., 2012; Kotak et al., 2013; Cundell et al., 2013; Enos et al., 2018; Keshri et al., 2020). PP2A is a trimeric holoenzyme complex consisting of ∼36 kDa catalytic (C subunit), a ∼65 kDa scaffold (A subunit) subunit, and a variable regulatory subunit (B subunit) that provides substrate specificity (reviewed in Barr et al., 2011; Gelens et al., 2018; Moura and Conde, 2019). Whether PP2A or phosphatases, in general, are involved in NE disassembly remains unknown.

Here, profiting from the highly stereotypical cell division of the one-cell *C. elegans* embryo, we discovered that B55^SUR-6^, an evolutionarily conserved regulatory subunit of PP2A-phosphatase, is essential for the timely nuclear envelope permeabilization (referred to as NEP), and the complete NE disassembly, marked by the complete loss of nuclear lamina, and several NPPs. In B55^SUR-6^ temperature-sensitive mutant embryos or B55^SUR-6^-depleted embryos, the lamina persists, and a subset of NPPs fail to disassemble during mitotic progression. This causes failure in merging of the parental genomes and improper segregation of sister chromatids. We reveal that the impact of B55^SUR-6^ on NEBD is not because of defective cell cycle progression or impaired localization of Aurora A or PLK-1. We further show that microtubules and dynein motor, which alone are dispensable for the onset of NEBD in *C. elegans* embryos, act synergistically with B55^SUR-6^ for efficient NE disassembly. Moreover, genetic epistasis and biochemical experiments indicate that the lamin LMN-1 is likely a direct target of B55^SUR-6^. In summary, our results indicate a vital but hitherto uncharacterized function of an evolutionarily conserved regulatory subunit of PP2A phosphatase B55^SUR-6^ in NEBD and suggest a model wherein a biochemical tug-of-war between kinases and PP2A phosphatase and the microtubules-dependent mechanical forces ensures timely NEBD in metazoans.

## Results

### The PP2A regulatory subunit B55^SUR-6^ facilitate NE disassembly during mitosis

In a candidate-based RNA interference (RNAi) screen to study the function of phosphoprotein phosphatase (PPP) family members during polarity establishment, we serendipitously noted that RNAi-mediated depletion of PP2A-regulatory subunit B55^SUR-6^ resulted in the appearance of multiple nuclei in the AB and P1 blastomeres (Figure S1A-S1C). This result is consistent with the previous notion that described the presence of more than one nucleus at the two-cell stage in B55^SUR-6^ temperature-sensitive mutant *or550* (referred to as *B55^SUR-6^ts*) embryos at the restrictive temperature (Figure S1A-S1C; O’Rourke et al., 2011). The occurrence of multiple nuclei in the AB/P1 blastomeres could be because of spindle assembly/maintenance defects that may result in chromosome segregation errors, and thus an extra nucleus or an inefficient NEBD, or a combination of both. To test these possibilities, we conducted the time-lapse recording of the one-cell embryos expressing the microtubule marker mCherry-tubulin or a kinetochore marker GFP-KNL-3, whose depletion has been associated with the kinetochore-null phenotype (Cheeseman et al., 2004; Kapoor and Kotak, 2019). Although chromosomes fail to congress on the metaphase plate in embryos depleted for B55^SUR-6^, we did not detect any significant change in microtubule nucleation capacity of the centrosomes or kinetochore localized KNL-3 levels in B55^SUR-6^-depleted embryos (Figure S1D-S1K). A similar conclusion was made for another kinetochore localized protein MEL-28, which is also a component of the NPC (Figure S1L-S1N; Galy et al., 2006). These data suggest that the formation of more than one nucleus in AB/P1 blastomeres upon B55^SUR-6^ depletion is likely not due to defective microtubule nucleation capacity of centrosomes or abnormal kinetochore organization.

In *C. elegans* early embryos, the permeability barrier of the nuclear envelope is breached at the onset of NEBD during prometaphase (referred to as NEP); however, the complete disassembly of the lamina occurs only ∼200 s later during metaphase-to-anaphase transition (Askjaer et al., 2002; Gorjanacz et al., 2007; Hachet et al., 2007; Portier et al., 2007). To analyze the role of B55^SUR-6^ in NEBD, we utilized the *C.elegans* strain coexpressing the single B-type lamin YFP-LMN-1 together with chromatin marker mCherry-HIS-58 (referred to as mCherry-H2B; Riemer et al., 1993; Liu et al., 2000). In control one-cell embryo, the female pronucleus is initially localized to the anterior after meiosis completion, and the male pronucleus is positioned at the posterior (Figure 1A and 1B). During prophase, the chromosomes condense while the female and male pronuclei migrate toward each other and meet at the embryo posterior. The two joined pronuclei then move to the cell centre and undergo NEBD, as monitored by the complete disappearance of the YFP-LMN-1 signal at the metaphase to anaphase transition (Figure 1A, 1B and 1D; Movie S1; reviewed in Cohen-fix and Askjaer, 2017; Pintard and Bowerman, 2019). Notably, the YFP-LMN-1 signal persists and never disappears during mitotic progression in a significant proportion (∼80%) of embryos depleted for B55^SUR-6^ (compare Figure 1C with 1B; Figure 1D; Movie S2). Quantification of the YFP-LMN-1 levels at the NE in control embryos suggested that the YFP-LMN-1 signal intensity progressively decreases ∼50 s before nuclear envelope permeabilization (NEP) (Figure 1E and 1F), which was followed by monitoring the diffusion of free nucleoplasmic mCherry-H2B signal out of the nucleus. In contrast, YFP-LMN-1 intensity is maintained in embryos depleted for B55^SUR-6^ (Figure 1E and 1F). Next, to examine the impact of B55^SUR-6^ depletion on the 3D architecture of the two pronuclei, we performed the volumetric analysis in control and B55^SUR-6^-depleted embryos (Figure 1G). As expected, the two pronuclei persisted upon B55^SUR-6^ depletion despite the steady decrease in the total nuclear volume (Figure 1H-1J).

**Figure 1.**
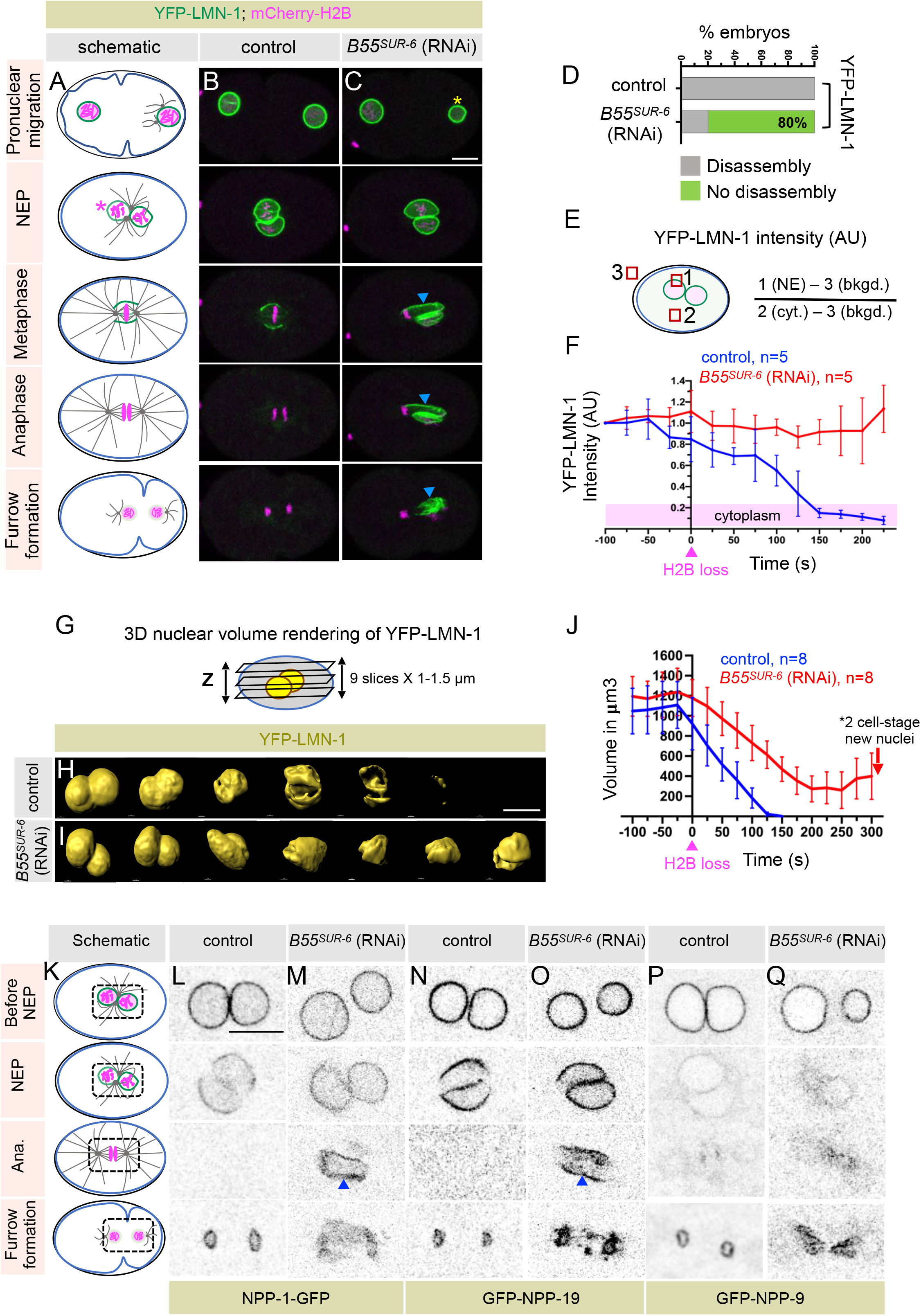
B55SUR-6 is critical for the lamina and nucleoporins disassembly during mitotic progression. (A-C) Schematics illustrate various stages of the cell cycle progression (A) and the corresponding images from the time-lapse confocal microscopy of *C. elegans* embryos coexpressing YFP-LMN-1 (green) and mCherry-H2B (magenta) in control (B) and *B55^SUR-6^ (RNAi)* (C). In this and subsequent images, embryos are approximately 50 μm in length, and the posterior is to the right. The magenta asterisk on the schematics represent nuclear envelope permeabilization stage (NEP) monitored by the expulsion of the nucleoplasmic mCherry-H2B signal from the nucleus. The yellow asterisk indicates the smaller size of the male pronucleus in *B55^SUR-6^ (RNAi)* embryos, and the blue arrowhead marks the persistence of LMN-1 signal during mitotic progression. The scale bar is 10 μm in this and other Figure panels. (D) Bar graph represents the percentage [%] of embryos that fail nuclear envelope disassembly during the metaphase-to-anaphase transition in *B55^SUR-6^ (RNAi)* compared to the control. [n=10 for control, n=30 for *B55^SUR-6^ (RNAi)*]. (E) Schematic representation of the quantification method used for analyzing YFP-LMN-1 fluorescence intensity on the nuclear envelope. bkgd., and cyt. represent background, and cytoplasmic intensity, respectively. (F) Quantification of YFP-LMN-1 fluorescence intensity on the nuclear envelope over time in control (blue) and *B55^SUR-6^ (RNAi)* (red) embryos as indicated in Figure panel E. Time ‘0’ is the nuclear envelope permeabilization stage (NEP) monitored by the expulsion of the nucleoplasmic mCherry-H2B signal from the nucleus. Error bars SD. (G) Schematic representation of image analyses method to measure the male and female pronuclei volume in control and *B55^SUR-6^ (RNAi)* embryos. Embryos were imaged with step size in *the z*-axis of 1 or 1.5 μm as indicated. Volume analysis was done using the surface 3D rendering feature in Imaris (Bitplane Inc; see Methods). (H, I) Representative 3D rendered sections of the male and female nuclei just before the onset of NEP from figures (B, C) depicting YFP-LMN-1 disassembly in control (H) in contrast to *B55^SUR-6^ (RNAi)* embryos (I). (J) Quantification of the volume (in μm^3^) for control and *B55^SUR-6^ (RNAi)* embryos as indicated in Figure panel G. Time ‘0’ is NEP, expulsion of the nucleoplasmic mCherry-H2B signal from the nucleus. Error bars SD. (K-Q) Schematics illustrate various stages of the cell cycle progression (K), and representative boxed sections in the schematics of the male and female pronuclei correspond to the images obtained from time-lapse confocal microscopy of embryos expressing NPP-1-GFP in control (L) and *B55^SUR-6^ (RNAi)* embryos (M); GFP-NPP-19 in control (N) and *B55^SUR-6^ (RNAi)* embryos (O); and GFP-NPP-9 in control (P) and *B55^SUR-6^ (RNAi)* embryos (Q). n=20, control; n=28, *B55^SUR-6^ (RNAi)* for NPP-1-GFP expressing embryos (27/28=96.43% proportion of RNAi embryos retain NPP-1-GFP signal upon *B55^SUR-6^* depletion); n=7, control; n=8, *B55^SUR-6^ (RNAi)* for GFP-NPP-19 expressing embryos [7/8; 87.5% proportion of RNAi embryos retain GFP-NPP-19 signal upon B55^SUR-6^ depletion]. In contrast, 93% of GFP-NPP-9 expressing embryos lose GFP-NPP-9 signal upon B55^SUR-6^ depletion (n=9, control; n=13, *B55^SUR-6^ (RNAi)*; 12/13; only 7% of embryos treated with RNAi show non-disassembly).

B55^SUR-6^ inactivation has been linked with the formation of the smaller size of male pronucleus in one-cell embryos (O’Rourke et al., 2011; Boudreau et al., 2019; Figure 1C). Thus, the persistence of nuclear lamina in the B55^SUR-6^-depleted embryos could be linked to defective male pronucleus. Therefore, we focused our attention on the female pronucleus and analyzed its growth after meiotic exit and its disassembly during mitotic progression in control and upon B55^SUR-6^ depletion (Figure S1O). The initial diameter and the growth kinetics of the female pronucleus in B55^SUR-6^-depleted embryos are comparable to the female pronucleus of control embryos (Figure S1P). Importantly, we found that the B55^SUR-6^ function in LMN-1 disassembly is not restricted to the male pronucleus but equally impacts the female pronucleus (Figure 1C; Movie S2). These results suggest that the role of B55^SUR-6^ in regulating male pronucleus size is distinct from its function in promoting NEBD.

We further sought to examine the role of B55^SUR-6^ in disassembling nucleoporin (NPPs) and inner nuclear envelope protein Emerin during NEBD. To this end, we first utilized strains that express tagged nucleoporin of the central channel (NPP-1-GFP), the inner ring (GFP-NPP-19), or the cytoplasmic filaments (GFP-NPP-9) (Figure S1Q; reviewed in Cohen-fix and Askjaer, 2017; Pintard and Bowerman, 2019). Analogous to LMN-1, the NE signal of NPP-1-GFP and GFP-NPP-19 persisted during metaphase-to-anaphase transition in 96.4% and 87.5% of the B55^SUR-6^ depleted embryos respectively, in contrast to 0% in the control embryos (Figure 1K-1O). Likewise, the NPP-1-GFP failed to disassemble in 100% of *B55^SUR-6^ts* mutant embryos at the restrictive temperature (Figure S5B). Interestingly, the cytoplasmic nucleoporin GFP-NPP-9 signal disappeared in 93% of the B55^SUR-6^-depleted embryos (Figure 1P and 1Q). To corroborate these findings with endogenous NPPs, we immunostained wild-type and *B55^SUR-6^ts* embryos with mAb414 antibodies, which detect NPPs with FG repeats (Aris and Blobel, 1989; Galy et al., 2003). Consistent with our analysis of NPP-1-GFP and GFP-NPP-19, we detected persistence of mAb414 signal during anaphase in *B55^SUR-6^ts* mutant embryos compared to wild-type embryos (data not shown). Next, we analyzed the role of B55^SUR-6^ in Emerin disassembly by examining the localization of mCherry-Emerin. Similar to the previous observations, we noted that the nuclear membrane undergoes a scission event between the two pronuclei, allowing the mingling of genomic contents of two paternal genomes in control embryos (Figure S1R and S1S; Rahman et al., 2015; Velez-Aguilera et al., 2020). However, this scission event failed to occur in a significant number of one-cell embryos depleted for B55^SUR-6^ (Figure S1T). Based on these results, we conclude that B55^SUR-6^ is essential for disassembling lamina, a subset of nucleoporins, and the inner nuclear envelope during mitotic progression.

### B55^SUR-6^ role in NEBD is not via regulating mitotic entry

How does B55^SUR6^ promote NEBD? Since CDK-1 and its associated cyclins are required for meiosis completion, mitotic entry, chromosome condensation, and NEBD (Boxem et al., 1999; van der Voet et al., 2009), one possibility is that B55^SUR-6^ is required for CDK-1 activation and, therefore, indirectly regulates NEBD. To test this assumption, we measured the interval between meiosis completion and pronuclear meeting in control and B55^SUR-6^-depleted embryos as a proxy for CDK-1 activation (Portier et al., 2007). The timing of meiosis completion to the pronuclear meeting in B55^SUR-6^-depleted embryos was similar to control [450s±61s in controls (n=8), versus 430s±59s in *B55^SUR-6^ (RNAi)* (n=10) embryos; p=0.491, ns; Figure 2A and 2B). To test this possibility more rigorously, we used a well-established image analysis method to quantify chromosome condensation in control and *B55^SUR-6^ (RNAi)* embryos expressing chromatin marker GFP-H2B (Figure S2A, and Methods; Maddox et al., 2006; Portier et al., 2007). The chromosome condensation kinetics of the female pronucleus in B55^SUR-6^-depleted embryos are identical to that of control embryos (Figure 2C and 2D). Though we did not observe any abnormality related to chromosome condensation or the growth of the female pronucleus upon B55^SUR-6^ depletion, there was a specific and significant delay of ∼120 s from the pronuclear meeting to NEP [120±31 s in control (n=11) versus 248±49s in *B55^SUR-6^ (RNAi)* (n=14); p<0.0001***; Figure 2A and 2B; Figure S1P]. These results suggest that B55^SUR-6^ regulates the initial stage of nuclear envelope permeabilization.

**Figure 2.**
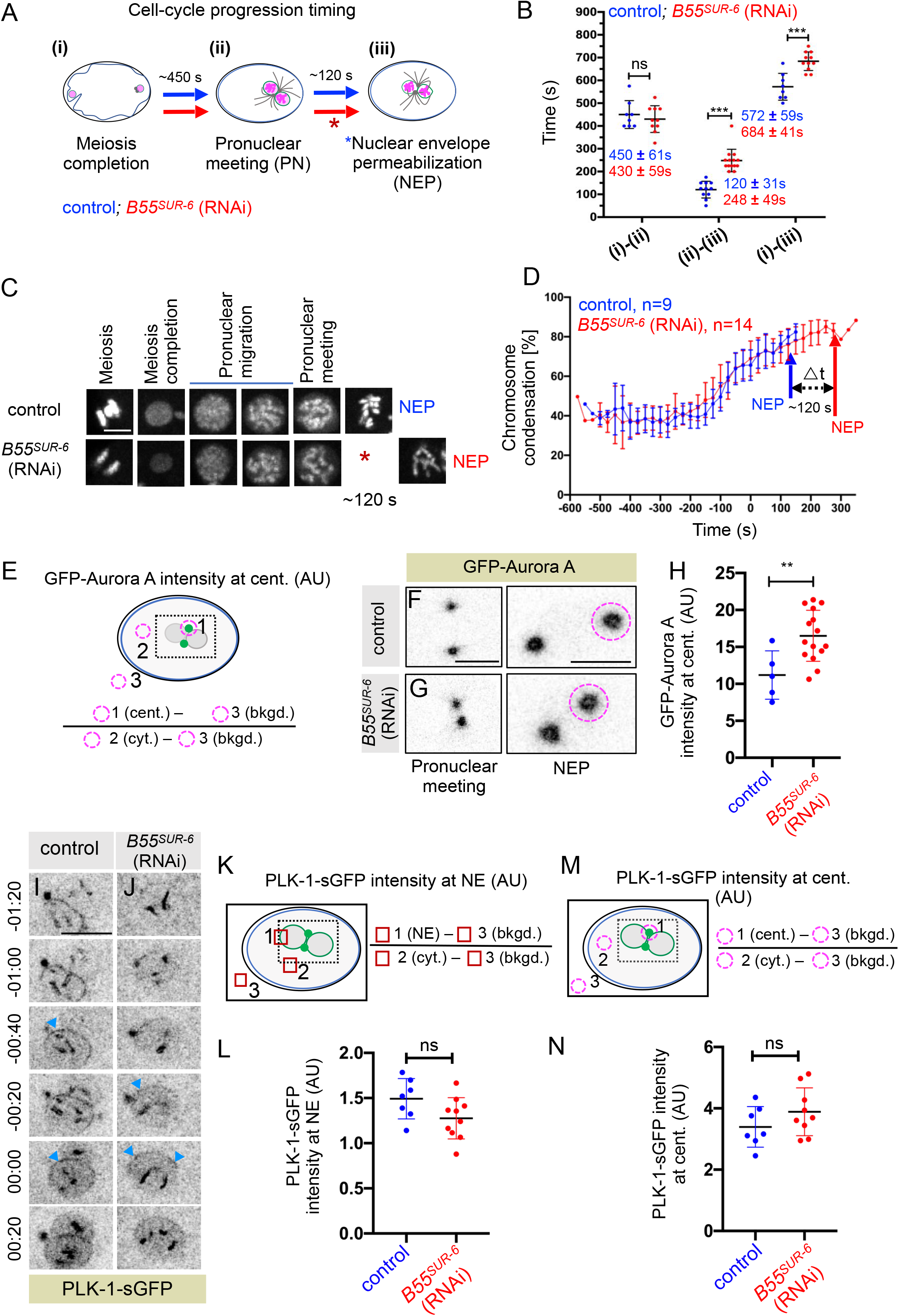
B55SUR-6 depletion delays NEP without affecting the mitotic entry. (A) Schematic representation of the cell cycle progression indicating various stages: meiosis completion (i), pronuclear meeting (ii), and NEP (iii). (B) Quantification of the cell cycle interval between the above-indicated stages (i), (ii), and (iii) in control (blue circle; n=8) and *B55^SUR-6^ (RNAi)* (red circle; n=11) embryos. The time interval between stages (i) and (ii) in control and *B55^SUR-6^ (RNAi)* embryos is ns [p=0.491]. However, there is a significant increase [p<0.0001] in a time interval between stages (ii) and (iii) of ∼120 s in *B55^SUR-6^ (RNAi)* (red circle; n=13) embryos when compared to control embryos (blue circle; n=11). Consequently, a similar delay persists from stage (i) to (iii). [(572±59 s, in control (blue circle, n=8) versus 684±41 s in *B55^SUR-6^ (RNAi)* (red circle, n=11) embryos [p<0.0001]. Error bars SD; p values were determined by two-tailed unpaired Student’s t-test. (C) Nuclear insets from maximum z-projections from time-lapse confocal microscopy of control and *B55^SUR-6^ (RNAi)* embryos expressing GFP-H2B at different stages of the cell cycle (C). Asterisk indicates that the NEP is delayed upon B55^SUR-6^ depletion by ∼120 s compared to the control. Scale bar 5 μm. (D) Quantification of chromosome condensation [%] of the female pronucleus in control (blue, n=9) and *B55^SUR-6^ (RNAi)* (red, n=14) embryos over time. The average value of condensation was calculated after aligning the embryos at the pronuclear meeting stage (t=0), which remains unchanged upon B55^SUR-6^ depletion (Figure 2B). See Supplementary Figure S2A for schematics of the method used for quantification; also see Methods for details. Error bars SD. (E) Schematic representation of the quantification method used to evaluate the GFP-Aurora A fluorescence intensity at the centrosome in control and *B55^SUR-6^ (RNAi)* embryos at the time of NEP. cent., bkgd., and cyt. represent centrosome, background, and cytoplasmic intensity, respectively. (F-H) Images from the time-lapse confocal microscopy of embryos expressing GFP-Aurora A in control (F) and *B55^SUR-6^ (RNAi)* (G) embryos. The dotted circle represents the area used for quantifying the GFP signal on the centrosome, shown as a scatter plot in (H). GFP-Aurora A levels at the centrosome in control (blue circle, n=5) and *B55^SUR-6^ (RNAi)* (red circle, n=15) embryos at NEP, monitored by the breakdown of the distinct contour of the nucleus in the DIC images (not shown). Scale bar 10 μm. Error bars SD; **\*\*,** P=0.0075 as determined by two-tailed unpaired Student’s t-test. (I, J) Images from the time-lapse confocal microscopy of embryos expressing PLK-1-sGFP in control (I) and *B55^SUR-6^ (RNAi)* (J) embryos. The blue arrowhead depicts the nuclear envelope. Note that the recruitment of PLK-1 to the nuclear envelope is unaffected in B55^SUR-6^-depleted embryos. Time stamps are relative to NEP (00.00 s), which was monitored by the breakdown of the distinct contour of the nucleus in the DIC images (not shown). Scale bar 10 μm. (K-N) Schematic representation of the quantification method used to evaluate the PLK-1-sGFP fluorescence intensity on the nuclear envelope (K) and on the centrosomes (M). Quantification of the fluorescence intensity of PLK-1-sGFP on the nuclear envelope (L) and the centrosome (N) in control (blue circle) and *B55^SUR-6^ (RNAi)* (red circle, n=10) depicted by the scatter plots. n=7 for control, and n>9 for *B55^SUR-6^ (RNAi)* embryos. cent., bkgd., and cyt. represent centrosome, background, and cytoplasmic intensity, respectively. Error bars SD; ns=0.070 (at NE) and p=0.202 (at centrosome), as determined by two-tailed unpaired Student’s t-test.

Earlier work has demonstrated that NEP occurs in two phases. In the first phase, the outermost components of NPCs are released, which allows the entry of molecules less than 40 nm in diameter into the nucleus. This is followed by the complete removal of NPCs, which allows the entry of larger particles of sizes significantly bigger than 40 nm (Terasaki et al., 2001; Lenart et al., 2003; Portier et al., 2007). To determine the function of B55^SUR-6^ on the kinetics of NEP, we examined the entry of Tetramethylrhodamine (TMR)-70 kDa dextran with a hydrodynamic radius of ∼36 nm as a marker for the first phase of permeabilization, and GFP-NMY-2 which has hydrodynamic radius >40 nm as a second phase of permeabilization (Citi and Kendrick-Jones, 1987; Lenart et al., 2003; Portier et al., 2007). In control embryos, the time interval between the entrance of GFP-NMY-2 and TMR-dextran is ∼45 s, which remained unchanged in embryos depleted of B55^SUR-6^ (Figure S2B-S2D), indicating that B55^SUR-6^ do not regulate the kinetics of NEP. Additionally, these results indicate that the impact of B55^SUR-6^ depletion on NEP and lamina/nucleoporins/inner nuclear membrane disassembly is not due to delayed mitotic onset or mitotic progression.

### B55^SUR-6^ orchestrates NEBD without influencing the localization and activity of Aurora A and PLK-1

In *C. elegans* one-cell embryo, the essential mitotic kinases Aurora A (also known as AIR-1 in *C. elegans*) and Polo-like kinase (PLK-1 in *C. elegans*) are implicated in NEBD (Portier et al., 2007; Hachet et al., 2007; Rahman *et al*., 2015; Martino et al., 2017; Velez-Aguilera et al., 2020; Nkoula et al., 2023). Therefore, we decided to explore whether B55^SUR-6^ could act upstream of Aurora A or PLK-1 to regulate NEBD. To this end, we monitored the levels of GFP-Aurora A at the centrosomes before nuclear envelope permeabilization in control, and B55^SUR-6^-depleted embryos. We did not detect any significant reduction in the intensity of the GFP-Aurora A signal at the centrosomes upon B55^SUR-6^ depletion, which is expected if B55^SUR-6^ would have been acting upstream of GFP-Aurora A in regulating NEBD (Figure 2E-2H). Also, the microtubule nucleation capacity of the centrosomes, which are partly regulated by Aurora A remains unchanged in B55^SUR-6^-depleted embryos (Figure S1D-S1G; Hannak et al., 2001; Kapoor and Kotak, 2019). It is worth noting that Aurora A depletion leads to ∼7 min delay in the NEP (Portier et al., 2007), in contrast to ∼2 min seen upon B55^SUR-6^ depletion (Figure 2B), indicating a distinct and Aurora A-independent mechanism of action of B55^SUR-6^ in regulating NEBD.

Transient accumulation of PLK-1 at the NE has been recently linked with NEBD (Martino et al., 2017; Linder et al., 2017). Therefore, we tested the possibility of B55^SUR-6^ involvement in enriching PLK-1 at the NE, thereby impacting NEBD. However, we did not observe any significant change in the fluorescence intensity of PLK-1-sGFP either at the NE or at the centrosomes in control and B55^SUR-6^-depleted embryos (Figure 2I-2N). This result is in line with phenotypic observations that RNAi-mediated depletion of B55^SUR-6^ in *B55^SUR-6^ts* embryos at restrictive temperature do not display any meiotic abnormality (data not shown), which is expected if B55^SUR-6^ would have been involved in controlling PLK-1 activity (Chase et al., 2000). These results suggest that B55^SUR-6^ has an essential and previously unidentified role in promoting timely NEBD that appears to be independent of its function in regulating the activity of CDK-1, Aurora A and PLK-1.

### B55^SUR-6^ promotes the proper merging of parental genomes

Since depletion or mutation in B55^SUR-6^ stabilizes nuclear lamina and a subset of NPPs, we assessed if the stabilization of nuclear lamina and NPPs is sufficient to prevent the merging of maternal and paternal genome in B55^SUR-6^-depleted embryos. For this purpose, we utilized embryos expressing photoconvertible H2B (Dendra-H2B), photoconverted the male pronucleus before pronuclei meeting, and followed the fate of the genomes after first mitosis in the two-cell stage as reported previously (Dendra-H2B) (Figure S2E; Rahman *et al*., 2015; Velez-Aguilera et al., 2020). The male and female genomes in control embryos, shown in magenta and green, are aligned on the metaphase plate. After segregation, these differently colored genomes are fully merged into a single nucleus at the two-cell stage (Figure S2F). In contrast, as expected, because of the improper NEBD, the male and the female genomes failed to merge in B55^SUR-6^-depleted embryos (Figure S2G). These data indicate that B55^SUR-6^ is critical for mixing paternal and maternal genomes.

#### Microtubule-dependent forces cooperate with B55^SUR-6^ for timely NEBD

Microtubules and NE-localized dynein/dynactin are propose to generate mechanical forces on the NE for efficient and robust NEBD in mammalian cells (Beaudouin et al., 2002; Salina et al., 2002). In the one-cell *C. elegans* embryo, when microtubules are depolymerized using Nocodazole or dynein motor is compromised, the male and female pronuclei fail to meet and undergo asynchronous NEP, which is followed by monitored expulsion of nucleoplasmic histone (H2B) signal out of the nucleus (Figure 3A and 3C; Movie S3). In this setting, the male pronucleus initiates permeabilization ∼70 s before the female pronucleus (referred to as Δt NEP; Figure 3D; Portier et al., 2007; Hachet et al., 2007; Hachet et al., 2012). Importantly, this asynchrony is independent of microtubules and dynein motor but dependent on the centrosomal pool of Aurora A kinase (Portier et al., 2007; Hachet et al., 2007). Since we uncovered that B55^SUR-6^ is essential for timely NEP and NEBD, we asked what happens to male and female pronuclei if the microtubules or dynein motor are compromised either in B55^SUR-6^-depleted or *B55^SUR-6^ts* embryos. Importantly, in ∼40% of the embryos that were depleted for B55^SUR-6^ and treated with Nocodazole, the Δt NEP between the male and female pronuclei was above 150 s (Figure 3E). Intriguingly, ∼15% of these embryos showed significantly extended asynchrony in NEP of more than 700 s (compare Figure 3B with 3A; Movie S4). Remarkably, we also noted that the chromosomes in the female pronucleus remained condensed for almost the entire duration of the time-lapse recording in these subset of embryos (Figure 3B, and data not shown). Identical findings were obtained for *B55^SUR-6^ts* embryos that were depleted for dynein motor at the restrictive temperature (Figure S3A-S3E).

**Figure 3.**
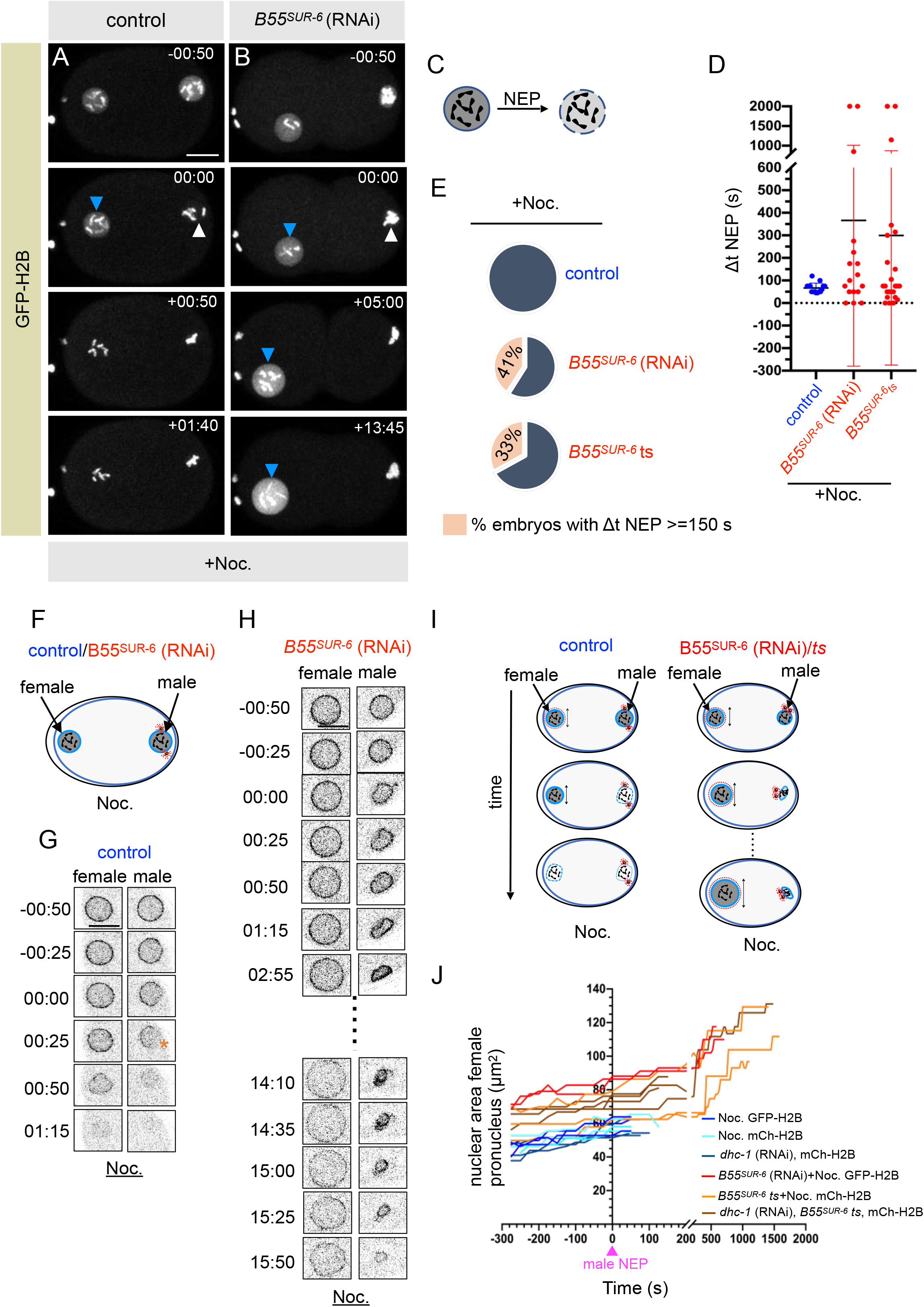
B55SUR-6 associated biochemical pathway cooperates with Microtubule-based mechanical forces for efficient NE disassembly. (A, B) Selected images from the time-lapse confocal microscopy of control embryos expressing GFP-H2B upon Nocodazole (Noc.) treatment (A) and *B55^SUR-6^ (RNAi)* embryos that are in addition treated with Noc. (B). Times are with respect to the expulsion of the nucleoplasmic GFP-H2B signal from the male pronucleus (white arrowhead). Please note that NEP of the female pronucleus (blue arrowhead) is significantly delayed in B55^SUR-6^-depleted embryos, which are also treated with Noc., in contrast to Noc. treated control embryos. The scale bar is 10 μm in this and other Figure panels. (C) Schematics of the loss of the nucleoplasmic H2B signal from the nucleus at the NEP. (D) Quantification of the interval (in s) between permeabilization of the male and female pronuclei (referred to as ΔNEP) in Noc. treated control (blue circles) or *B55^SUR-6^ (RNAi)/ B55^SUR-6^ts* (red circles) embryos that are treated with Noc. Note an exaggerated ΔNEP in *B55^SUR-6^ (RNAi)/B55^SUR-6^ts* + Noc. treated embryos. Also note the tight window of data points from the pooled set of control embryos (blue circles) with an average ΔNEP of ∼65 s (66±22 s, n=15) in contrast to a wide spread of data points in B55^SUR-6^-depleted (n=17; red circles)/*B55^SUR-6^ts* (n=24; red circles) embryos that are also treated with Noc. Error bars SD. (E) Pie-chart depicting the pooled percentage of embryos showing asynchronous NEP (ΔNEP) in Noc.-treated control, *B55^SUR-6^ (RNAi)*, or *B55^SUR-6^ts* embryos expressing GFP-H2B or mCherry-H2B and are in addition treated with Noc. Note that ∼41% *B55^SUR-6^ (RNAi)* and ∼33% *B55^SUR-6^ts* embryos that are treated with Noc. show significantly longer asynchronous NEP (>= 150) between male and female pronucleus than Noc. treated embryos alone. (n=7/17 in *B55^SUR-6^ (RNAi)* and n=8/24 in *B55^SUR-6^ts*). (F) Schematic illustration of control and *B55^SUR-6^ (RNAi)* embryos treated with Noc. showing the effect of the Noc. and the relative positions of the nuclei (F). (G, H) Male and female nuclear insets from the embryos expressing NPP-1-GFP that are either treated with Noc. (G) or *B55^SUR-6^ (RNAi)* embryos that are also treated with Noc. (H). Note the excessive delay in nuclear-shape deformation in the female pronucleus in *B55^SUR-6^ (RNAi)*+Noc.-treated embryos compared to the male pronucleus or the female pronucleus of control Noc.-treated embryos till the movie was recorded. (I) Schematic representation to analyze the asynchrony in the female pronuclear size of control and B55^SUR-6^-depleted embryos when these embryos are compromised for microtubules by Noc. or impaired for dynein motor function by *dhc-1(RNAi)*. (J) Individual trajectories of the female nuclear size (in μm^2^) from selected control or *B55^SUR-6^* (RNAi)/*B55^SUR-6^ts* embryos expressing either GFP-H2B or mCherry-H2B, which are in addition treated with Noc. or depleted for dynein motor by *dhc-1 (RNAi)* as indicated. The method to quantify the nuclear size is described in Methods. Nuclear expansion trajectories of individual female pronucleus in these conditions embryos are aligned with respect to male NEP (time ‘0’). Y-axis is the nuclear size area in μm^2^.

Since NEP is closely followed by the collapse of the spherical shape of the pronuclei, we next sought to dissect the nuclear envelope morphology in these experimental settings. To this end, we performed time-lapse confocal imaging of control or B55^SUR-6^-depleted embryos expressing NPP-1-GFP that were in addition treated with Nocodazole. In the control embryos treated with Nocodazole, the male and female pronuclei are initially delineated by a homogenous fluorescence intensity of NPP-1-GFP. The nuclear shape of the male pronucleus is crumpled ∼70 s before the female pronucleus, followed by the complete disappearance of the NPP-1-GFP signal at the NE (Figure 3F and 3G). In B55^SUR-6^-depleted embryos with compromised microtubules, the male pronucleus shows crumpling of the nuclear envelope, however as expected, the NPP-1-GFP signal persisted given our observation on the role of B55^SUR-6^ in complete disassembly of the NE (Figure 3F and 3H). Strikingly, the female pronucleus that does not carry centrosomes and is far away from the male pronucleus shows a remarkably roundish nuclear envelope that lasts for more than 150 s in ∼40% of the embryos (Figure 3C, 3F, and 3H). And in 15% of these embryos, the female pronucleus stays roundish and continues to grow for a substantially longer duration of ∼700 s or more (Figure 3H, 3I and 3J).

Similarly, *B55^SUR-6^ts* embryos that are depleted for dynein motor show a uniform and roundish morphology of the female pronucleus for a significantly longer duration than the dynein-depleted control embryos (compare Figure S3B with S3A; Movies S5 and S6). These data suggest that while microtubule-dependent mechanical forces alone are insufficient for NEBD in *C. elegans* zygote, and that these forces act redundantly with the B55^SUR-6^-dependent NE disassembly pathway for efficient and robust NEBD program (see discussion).

### B55^SUR-6^ is robustly localized to the nucleus at NEP

To analyze how B55^SUR-6^ helps in NEBD, we investigated the localization of B55^SUR-6^ during mitotic progression. To this end, we utilized the *C. elegans* strain, coexpressing endogenous-tagged B55^SUR-6^ with GFP and FLAG-tag (hereafter referred to as GFP-B55^SUR-6^; Bel Borja et al., 2020) and mCherry-tubulin. Time-lapse imaging of this line revealed that GFP-B55^SUR-6^ is localized to the nucleus above its cytoplasmic signal in the prophase before NEP (Figure S4A). The GFP-B55^SUR-6^ levels significantly rises in the nucleus concomitantly with the accumulation of free mCherry-tubulin signal during NEP (Figure S4A-S4C; Movie S7). Interestingly, a previous work has reported that the GFP-tagged catalytical subunit of PP2A-LET-92 is robustly localized to the nucleus at or a few seconds before NEP (Schlaitz et al., 2007). Altogether these localization studies suggest that the assembly of the PP2A-B55^SUR-6^ complex inside the nucleus could play an essential function for the efficient NEBD (see discussion).

Next, to test if the nuclear enrichment of GFP-B55^SUR-6^ either at or a few seconds before NE permeabilization could be linked with NE disassembly, we analyzed the localization of GFP-B55^SUR-6^ in embryos in Nocodazole-treated conditions where male and female pronuclei undergoes asynchronous NEP with Δt of ∼70 s (Figure 3). As expected, the GFP-B55^SUR-6^ signal is present in the nucleus during prophase. However, the GFP-B55^SUR-6^ fluorescence intensity is significantly enriched inside the male pronucleus ∼50 s before the NEP of the female pronucleus (Figure S4D and S4E; Movie S8). Altogether, these observations raise a tempting possibility that the nuclear enrichment of the PP2A-B55^SUR-6^ complex promotes efficient NE permeabilization and its disassembly.

### LMN-1 is a potential downstream target of B55^SUR-6^

Hyper-phosphorylation of the lamina and nucleoporins by kinases results in disassembly of the NE during mitosis (Gerace and Blobel, 1980; Heald and McKeon, 1990; Muhlhauser and Kutay, 2007; Laurell et al., 2011; Mehsen et al., 2018; Velez-Aguilera et al., 2020; reviewed in Machowska et al., 2015). It has been proposed that efficient nuclear disassembly might require the removal of the interphase pattern of phosphorylation from the lamina and nucleoporins for efficient NEBD (reviewed in Machowska et al., 2015). However, to the best of our knowledge, no report shows if this occurs *in vivo*. We reasoned if PP2A-B55^SUR-6^ functions as a critical phosphatase required to remove certain inhibitory phosphate group/s from NPP/s or LMN-1 for NEBD, then the simultaneous depletion of B55^SUR-6^ with that particular NPP/s or LMN-1 should dissolve nuclear envelope that persists in B55^SUR-6^ depleted embryos (Figure S5A). To explore this possibility, we performed a candidate-based RNAi suppressor screen by depleting B55^SUR-6^ along with various NPPs or a single B-type lamin LMN-1 (Liu et al., 2000). RNAi-mediated depletion of 22 NPPs out of a total of 27 encoded by the *C. elegans* genome (reviewed in Cohen-fix and Askjaer, 2017; Pintard and Bowerman, 2019) failed to disassemble the NPP-1-GFP signal seen in B55^SUR-6^-depleted embryos (Supplementary Table 1).

Next, we assessed the impact of LMN-1 depletion on NEBD in either control embryos expressing NPP-1-GFP or *B55^SUR-6^ts* embryos expressing NPP-1-GFP. LMN-1 depletion impacts the nuclear morphology, as indicated previously (Figure 4A; Liu et al., 2000); however, LMN-1 depletion had no impact on NE dissolution during mitotic progression (100% of embryos depleted of LMN-1 dissolve their NE during mitotic progression; Figure 4A). Notably, in stark contrast with *B55^SUR-6^ts* embryos, where the NPP-1-GFP signal persists in 100% of embryos (Figure 4B; Figure S5B), depletion of LMN-1 in *B55^SUR-6^ts* embryos resulted in the complete disappearance of the NPP-1-GFP nuclear envelope signal in ∼80% of the embryos (compare Figure 4C with 4B; Figure S5B). We further validated these findings in *C. elegans* strain expressing another nucleoporin marker GFP-NPP-19 in the B55^SUR-6^-depleted conditions (Figure S5B). Since the persistent nuclei observed in *B55^SUR-6^ts* embryos disassembled upon inactivation of LMN-1, we conclude that the lamina acts downstream of B55^SUR-6^ for timely NE dissolution.

**Figure 4.**
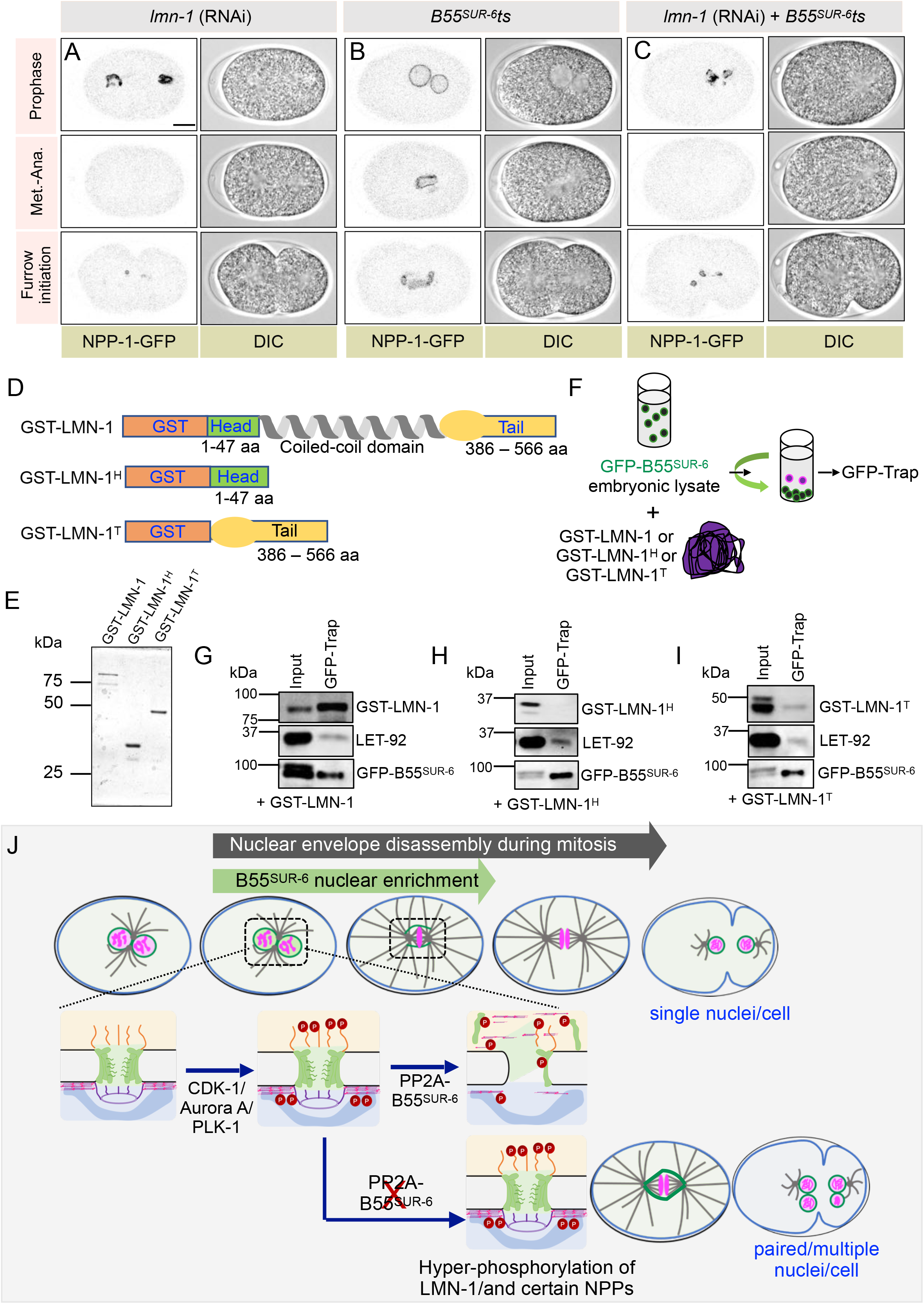
LMN-1 is a bona-fide target of B55^SUR-6^ for NEBD. (A-C) Images from time-lapse confocal microscopy in combination with DIC microscopy at various cell cycle stages as indicated in embryos expressing NPP-1-GFP in *lmn-1 (RNAi)* (A) in *B55^SUR-6^ts* background (B) or *lmn-1 (RNAi)* in *B55^SUR-6^ts* background (C). 100% of LMN-1-depleted embryos show complete disassembly of NPP-1-GFP signal (middle panel), as opposed to 0% in *B55^SUR-6^ts* embryos. Note that the majority (80%) of the embryos depleted of LMN-1 in *B55^SUR-6^ts* mutant embryos show disassembly of NPP-1-GFP signal [n=29, *B55 ^SUR-6^ ts* mutant embryos; n=7, *lmn-1* (RNAi) embryos, and n=10, *lmn-1 (RNAi)*, *B55^SUR-6^ts* mutant embryos]. (D) Schematics of Glutathione S-transferase (GST)-tag LMN-1 (GST-LMN-1) or GST-tagged head-domain (1-47 aa) of LMN-1 (GST-LMN-1^H^), or GST-tagged tail domain (386-566 aa) of LMN-1 (GST-LMN-1^T^). (E) Coomassie-stained gel loaded with recombinant GST-LMN-1, GST-LMN-1^H^, and GST-LMN-1^T^. (F) Schematic representation of the Co-immunoprecipitation (IP) by GFP-Trap using embryonic lysate from embryos expressing endogenously-tagged B55^SUR-6^ with the GFP and the FLAG tag (referred to as GFP-B55^SUR-6^) after incubation with GST-LMN-1 or GST-LMN-1^H^ or GST-LMN-1^T^ at room temperature for 15 min. (G-I) IP by GFP-Trap from embryonic lysates expressing GFP-B55^SUR-6^ that were incubated with GST-LMN-1 (G) or GST-LMN-1^H^ (H) or GST-LMN-1^T^ (I) as indicated in Figure panel (F). Resulting blots were probed for GST (to detect GST-tagged recombinant LMN-1-full length, or GST-LMN-1H, or GST-LMN-1T), LET-92 (the catalytic subunit), and FLAG (to detect GFP-B55^SUR-6^). Input (2% of total), GFP-Trap fraction-33% of the total. Please note that 3% of IP fraction was loaded for the FLAG detection. (J) Model for PP2A-B55^SUR-6^-dependent nuclear disassembly pathway at the first-embryonic division of *C. elegans.* In control embryos, the presence, and the rapid import of PP2A-B55^SUR-6^ at NEP (green) in nucleoplasm may help erasing the inhibitory phosphorylation marks on the nuclear envelope components, including the lamina LMN-1, and NPPs. In addition, efficient NEBD depends on microtubule-dependent forces. In the absence of PP2A-B55^SUR-6^, hyperphosphorylation of some of the key residues prevents timely NEP and NEBD and, thus, leads to chromosome missegregation and the failure of merging of the maternal and paternal genomes.

To determine if LMN-1 could be a direct target of B55^SUR-6^, we generated recombinant full-length LMN-1 with Glutathione S-transferase (GST) tag in *E*. *coli* (Figure 5D and 5E). Subsequently, we performed biochemical interaction studies by mixing GST-LMN-1 with the embryonic lysates made from *C. elegans* strains expressing GFP-B55^SUR-6^ (Figure 5F). Co-immunoprecipitation (IP) with anti-GFP nanobody coupled to agarose beads revealed that GFP-B55^SUR-6^ efficiently interacts with GST-LMN-1, and PP2A catalytical subunit LET-92 (Figure 5G). Since LMN-1 disassembly is regulated by phosphorylation events at the LMN-1 head (1-47 aa) or tail (386-566 aa) domains (Velez-Aguilera et al., 2020; reviewed in Machowska et al., 2015), we further generated recombinant LMN-1 head and tail domain fused to GST to analyze their interaction with GFP-B55^SUR-6^ (Figure 5D-5F). This analysis revealed that the C-terminal tail (GST-LMN-1^T^), but not the head domain (GST-LMN-1^H^), precipitated with the GFP-B55^SUR-6^ (compare Figure 5I with 5H). These results suggest that GFP-B55^SUR-6^ interacts with the sole lamina component LMN-1 in *C. elegans* embryos and allude to an exciting possibility that PP2A-B55^SUR-6^ complex function might be associated with dephosphorylation of some of the critical phosphorylated residues at the C-terminal region of the lamina for timely NE disassembly, and thus for allowing the merging of paternal and maternal genomes (see discussion).

## Discussion

Disassembly of the NE is critical for spindle assembly and chromosome segregation in animal cells. How cells efficiently disassemble the NE comprising of large complexes such as NPCs, lamina, and nucleoskeleton and cytoskeleton (LINC) linker during mitotic progression remains incompletely understood. Previous work established that mitotic kinases are the major drivers of the NPCs and lamina disassembly (reviewed in Cohen-fix and Askjaer, 2017; Pintard and Bowerman, 2019; Ungricht and Kutay, 2017; Kutay et al., 2021). These kinases hyperphosphorylate various nucleoporins (NPPs) and lamin and thus promote NEBD. Using *C. elegans* one-cell embryos that provide developmental context to characterize the mechanisms of NEBD, we unexpectedly discovered that an evolutionarily conserved regulatory subunit of PP2A, B55^SUR-6^ is a key for NEP and the complete dissolution of lamin and NPCs during mitotic progression (Figure 4J). This function of B55^SUR-6^ is crucial for proper chromosome segregation and the mingling of the parental genomes.

### B55^SUR-6^ is a crucial regulator of NEBD

B55^SUR-6^ promotes NEBD by facilitating lamin and a subset of nucleoporins disassembly. The failure of NEBD in B55^SUR-6^-depleted one-cell embryos leads to the formation of more than one nucleus in the AB or P1 blastomeres or both at the two-cell stage (this work, and O’Rourke et al., 2011). This is most likely because of 1) inefficient chromosomes capturing by the microtubules because of the persistence of the nuclear structure that creates a barrier and causes the random distribution of the genetic material during anaphase and the formation of extra-nuclei, and 2) the failure in the pronuclear scission leading to paired-nuclei formation. In many B55^SUR-6^-depleted embryos, the chromosomes fail to align on the metaphase plate, supporting the first hypothesis. The inefficient chromosome capturing by the microtubules is seemingly not because of defective kinetochore organization or the microtubule emanation capacity of the centrosomes. In addition, in a subset of embryos, we also noted the absence of pronuclear scission and, thus, paired nuclei formation. Altogether, these observations argue in favour of both possibilities, which results in more than one nucleus at the two-cell stage.

How does B55^SUR-6^ regulate NEBD? A plethora of work in the past has established that mitotic kinases such as CDK-1, Aurora A, and PLK-1 facilitate NEBD in *C. elegans* embryos (Boxem et al., 1999; van der Voet et al., 2009; Portier et al., 2007; Hachet et al., 2007; Hachet et al., 2012; Rahman et al., 2015; Martino et al., 2017; Velez-Aguilera et al., 2020; Nkoula et al., 2023). Therefore, one plausibility was that B55^SUR-6^ indirectly promotes NEBD by controlling the localization/activity of these kinases. However, the interval between meiosis completion to pseudocleavage regression/pronuclei meeting, which serves as phenotypic markers of the mitotic entry, remains unaltered upon B55^SUR-6^ depletion. These data indicate that the mitotic onset, which chiefly relies on global CDK-1 activation, remains unaffected in B55^SUR-6^-depleted embryos. Likewise, the localization of Aurora A and PLK-1 is largely unperturbed in B55^SUR-6^-depleted embryos, suggesting that B55^SUR-6^ role in NEBD is not via regulating the localization of these kinases. While we could not directly assess the PLK-1 activity upon B55^SUR-6^ depletion, however, given our observation that B55^SUR-6^ depletion does not phenocopy PLK-1 loss with respect to NEBD during meiosis and cell cycle progression (Chase et al., 2000; Noatynska et al., 2010; Cabral et al., 2019), we think that B55^SUR-6^ do not regulate PLK-1 activity. The genetic analysis reveals that B55^SUR-6^ acts upstream of lamina since the depletion of LMN-1 in either *B55^SUR-6^ts* or B55^SUR-6^-depleted embryos dissolves NE. Furthermore, the biochemical interaction analysis revealed that B55^SUR-6^ in complex with its catalytical subunit LET-92 interacts with the globular tail domain (386-566 aa) of LMN-1. Since phosphorylation of the head (1-47 aa) and the tail domain of lamin promote its disassembly during mitosis (Velez-Aguilera et al., 2020), we speculate that dephosphorylation of certain phosphorylated residues of LMN-1 by the PP2A-B55^SUR-6^ complex could be a decisive step in timely NEBD, at least in the context of *C. elegans* zygote (Figure 4J). In B55^SUR-6^ depleted condition, the LMN-1 would remain phosphorylated at those residues, creating a roadblock for NE disassembly (Figure 4J). Though our genetic data suggest that nucleoporins do not epistatically interact with B55^SUR-6^, at this moment, we cannot firmly rule out a possibility that more than one nucleoporin/s could be a biochemical target of PP2A-B55^SUR-6^ for efficient NEBD (Figure 4J). A key challenge for future work would be identifying the target/s and the phosphorylated marks that PP2A-B55^SUR-6^ dephosphorylates to orchestrate NEBD.

### Microtubule, and dynein motor-based mechanical forces cooperate with **B55^SUR-6^ to initiate NEBD**

In mammalian somatic cells, microtubules and dynein motor-mediated mechanical forces are implicated in tearing down the NE at the initial phase of NEBD (Beaudouin et al., 2002; Salina et al., 2002). Interestingly, centrosomally nucleated microtubules and the dynein motor alone are dispensable for NEBD onset. This was beautifully demonstrated in conditions where male and female pronuclei failed to meet because of reduced microtubule nucleation or dynein motor dysfunction (Hachet et al., 2007; Portier et al., 2007). In such experiments, the male and female pronucleus undergoes asynchronous NEBD. The male pronucleus carrying the centrosome commences NEP ∼70 s before the female pronucleus (this study; Hachet et al., 2007). The NEP asynchrony between male and female pronucleus was shown to be microtubules and dynein motor-independent but Aurora A-dependent, which is enriched at the centrosomes, and thus present in the vicinity of the male pronucleus (Portier et al., 2007; Hachet et al., 2007). Here, by treating B55^SUR-6^-depleted embryos with microtubule poison Nocodazole, we unexpectedly observed an exaggerated increase in NEP asynchrony between male and female pronucleus. Intriguingly, in a subset (∼15%) of embryos, the female pronucleus failed to undergo NEP for more than ∼700 s, in contrast to the ∼70 s in Nocodazole-treated control embryos. In such embryos, the female pronucleus retains all the chromosomes in a condensed state and steadily increases in size. These data suggest that initial NEP and, subsequently, NEBD are critical events for DNA decondensation and act as a timer to reset the nuclear size and volume. Similar observations were made in embryos depleted for dynein motor in *B55^SUR-6^ts* mutant embryos. While not all B55^SUR-6^-depleted and microtubule/dynein compromised embryos show this exaggerated asynchrony in NEP, we are of the opinion that incomplete RNAi efficiency and residual microtubules may account for this observation. Overall, this work, and the recent work from Velez-Aguilera et al., which implicated the function dynein-dependent cortical pulling forces in the pronuclear envelopes scission during metaphase to anaphase transition (Velez-Aguilera et al., 2022) imply that mechanical forces, which alone are dispensable for NEBD in *C. elegans* embryos, act synergistically with cellular biochemical machinery for efficient NE disassembly.

### PP2A-B55^SUR-6^: a missing link for NEBD during embryogenesis

Lamina depolymerization’s kinetics and timing vary across different species. For instance, in human somatic cells, NEBD occurs in prophase (reviewed in Hetzer, 2010; Ungricht and Kutay, 2017). In contrast, in *C. elegans* and Drosophila embryos, the lamina stay partly intact during prophase and only completely dismantled during the metaphase to anaphase transition (Paddy et al., 1996; Lee et al., 2000; Askjaer et al., 2002; Hachet et al., 2007; Portier et al., 2007; Katsani et al., 2008 Hachet et al., 2012). We uncovered that B55^SUR-6^ is crucial for initial NEP. In embryos depleted for B55^SUR-6^, the NEP is significantly delayed for ∼120 s, which is ∼50% of the time that embryos spent from NEP to anaphase onset. Remarkably, NEP in *C. elegans* embryos coincides with the downfall in cyclin B levels and possibly with a decrease in CDK-1 activity that occurs ∼200 s before metaphase to anaphase transition (Kim, Lara-Gonzalez et al., 2017). Since CDK-1/cyclinB-mediated phosphorylation is linked to cell-cycle-dependent regulation of PP2A-B55 in human cells (Schmitz et al., 2010; Cundell et al., 2016; reviewed in Nilsson, 2018), we hypothesize that decrease in CDK-1 activity at the NEP would be a decisive step in controlling PP2A-B55^SUR-6^ activity, and thus for the initiation of NEP and complete disassembly of the NE. Notably, B55^SUR-6^ and its catalytic subunit LET-92 are enriched in the nucleoplasm (this work, and Schlaitz et al., 2007). Therefore, we envisage that PP2A-B55^SUR-6^ spatial localization in the one-cell embryos makes this complex competent to carry out time-dependent NE disassembly in *C. elegans* and possibly in other embryos. In summary, our findings link the role of the PP2A-B55^SUR-6^ complex in NE disassembly and suggest the possibility of a biochemical cross-talk between kinases and phosphatase/s to choreograph NEBD.

## Materials and Methods

### *C. elegans* Strains and drug treatment

*C. elegans* wild-type (N2) and transgenic lines expressing GFP/sGFP/mCherry-tagged proteins were maintained at either 20°C or 24°C. Temperature-sensitive strains were maintained at 15°C. A list of all strains used in the study has been provided in Supplementary Table 2.

Microtubule poison Nocodazole treatment was performed as described (Bienkowska and Cowan, 2012; Kapoor and Kotak, 2019). In brief, worms were dissected in Nocodazole (10 μg/ml; Sigma Aldrich: M1404) containing egg buffer (118 mM NaCl, 40 mM KCl, 3.4 mM MgCl2, 3.4 mM CaCl2 and 5 mM HEPES pH 7.4; also see Boyd et al., 1996) and the drug could enter in the embryos because of the permeability of the eggshell during meiosis II (Johnston et al., 2006). The efficiency of Nocodazole was determined by the inability of the male and female pronuclei to migrate.

### RNAi experiments

Bacterial RNAi feeding strains for *B55^SUR-6^*, *lmn-1*, *dhc-1,* and several nucleoporins (NPPs) (Supplementary Table 2) were obtained from the *C. elegans* ORFeome RNAi library (Rual et al., 2004) or Source BioScience (Kamath et al., 2003). Feeding vectors for a few NPPs (Supplementary Table 1) were generated by amplifying the gene from either cDNA/gDNA and cloning in L4440 feeding vector (list of all NPPs used in screen provided in Supplementary Table 1). RNAi against all the genes (Supplementary Table 2) used in this work was performed by feeding animals starting at the L2 or L3 stage with bacteria expressing the corresponding dsRNAs at 20°C or 24°C for 24-48 h before analysis. While performing double depletion using RNAi, the single depletion was performed side by side to observe the single RNAi-mediated phenotype.

RNAi experiments in *B55^SUR-6^ts* mutant strain (EU1062) was performed by picking L3-L4 on RNAi feeding plate for 12-20 h at 15°C followed by shifting the feeding plates to the restrictive temperature of 25°C for 12-24 h, as in *dhc-1 (RNAi)* or *lmn-1 (RNAi)*.

### Time-lapse microscopy

For recording embryos, gravid worms were dissected in M9 or egg buffer and transferred onto a 2% agarose pad containing slides using a mouth pipette. These were then covered with a 20 X 20 mm coverslip. Time-lapse Differential Interference Contrast (DIC) microscopy, Confocal microscopy in combination with DIC were performed on such embryos either on IX53 (Olympus Corporation, Japan) with Qimaging Micropublisher 5.0 Colour CCD Camera (Qimaging, Canada) with 100X 1.4 NA objective, or FV3000 Confocal system with high-sensitivity cooled GaAsP detection unit (Olympus Corporation, Japan) using 60X 1.4 NA objective. Images were collected at intervals of 5-25 s per frame. Movies were subsequently processed using ImageJ maintaining relative image intensities within a series. Z-stack series were projected as maximum intensity projections for embryos expressing mCherry-tubulin, GFP-Aurora A, mCherry-H2B, GFP-KNL-3, GFP-MEL-28, and Dendra-H2B.

### Indirect immunofluorescence

Embryo fixation and staining for indirect immunofluorescence was performed mostly as described (Gönczy et al., 1999), using 1:200 mouse anti-α-tubulin antibodies (DM1A, Sigma), in combination with 1:200 rabbit anti-mAB414 (Biolegend: 902907). Embryos were fixed in methanol at –20°C for 30 minutes and incubated with primary antibodies for one hour at room temperature. Secondary antibodies were Alexa488-coupled anti-mouse and Alexa568-coupled anti-rabbit, both used at 1:500. Confocal images were acquired on a FV3000 Confocal system with high-sensitivity cooled GaAsP detection unit (Olympus Corporation, Japan) using 60X objective with NA 1.4 oil and processed in ImageJ and Adobe Photoshop, maintaining relative image intensities.

### Photoconversion experiment

Photoconversion experiments of worms expressing photoconvertible histone, Dendra2-H2B (strain OCF69) were performed as described in (Rosu and Cohen-Fix, 2017). A circular ROI around the male pronucleus was illuminated with a 405 nm laser set at low power of 0.9% for 20 ms and pixel dwell time of 10 ms. A green-to-red photoconversion of the labeled H2B chromosomes was observed, and the embryos were recorded using z-stacking with 1.5 μm with 25 s time interval as described before. Image stacks were processed using Fiji and are represented as maximum z-projections.

## Data analysis

Fiji/ImageJ (https://fiji.sc/), Imaris (Bitplane Inc.), and GraphPad Prism were used to perform quantitative analysis. Fiji was used to quantify nuclear area. Imaris was used for 3D rendering of the nucleus and quantification of volume. The p-value was considered to be significant if p < 0.05 using GraphPad Prism 8. The p values are either mentioned in text or on the graph and the significance values are mentioned as n.s., >0.05; *p<0.05; **p<0.01; ***p<0.001.

### Assigning time ‘0’

For most data sets and graphs, t=0 corresponds to nuclear envelope permeabilization (NEP). This is the frame at which there is a visible exclusion of nucleoplasmic H2B signal or inclusion of tubulin signal. However, in the context of chromosome condensation kinetics, nuclear size expansion-related graphs the pronuclear meeting timing is considered ‘0’ as indicated.

### Quantification of LMN-1 fluorescence intensity on the NE

A fixed rectangular ROI (ROI of 2.4 μm2) was placed on the nuclear envelope at the time intervals indicated on the graph on single z-sections of the image. The integrated fluorescence intensity in this ROI and a similarly sized ROI in the cytoplasm and background were determined using Fiji/ImageJ. Fluorescence intensity at the Nuclear envelope and cytoplasm was subtracted from the background, and the ratio of NE/cytoplasm to indicate NE enrichment was used for plotting the graph. Signal intensities were compared using a two-sample unpaired t-test in GraphPad Prism.

### Quantification of mCherry and GFP fluorescence intensity in mCherry-tubulin, GFP-AIR-1 and PLK-1-sGFP expressing embryos

Centrosome intensity of mCherry-tubulin (ROI of area 12.574 μm^2^), GFP-AIR-1 (ROI of area 28.67 μm^2^), and PLK-1-sGFP (ROI of area 3.4 μm^2^) expressing embryos was quantified by taking the ratios of fluorescent intensities at the centrosome w.r.t cytoplasm and corrected for the background. The nuclear envelope intensity of PLK-1-sGFP (ROI of area 2.96 μm^2^) expressing embryos was also quantified with the method described above for LMN-1.

### Quantification of KNL-1 and MEL-28 intensity at the kinetochore

Quantification of KNL-1 and MEL-28 kinetochore localization in GFP-KNL-1 and GFP-MEL-28 expressing embryos at the metaphase stage of one-cell embryos was performed on maximum intensity projections show in schematic in Figure S1J, as in Dumont et al., 2010. In brief, a rectangular ROI was drawn around the kinetochore signal, and integrated pixel intensity was measured. The ROI was expanded by 5 pixels and the difference in the integrated intensity between this ROI and the original ROI was used to define the background. Integrated kinetochore fluorescence is calculated as the value of the original ROI after background subtraction. For data representation on plot, the values have been normalized to 1 with respect to the maximum kinetochore intensity observed in control values.

### Quantification of nuclear fluorescence intensity for GFP-B55^SUR-6^ and mCherry-tubulin nuclear enrichment

A fixed circular ROI (area 3.4 μm^2^) was placed manually inside the nucleus at time intervals indicated on the graph. The integrated fluorescence intensity in this ROI and a similarly sized ROI in the cytoplasm and background were determined using Fiji/ImageJ. Fluorescence intensity at the centrosome and cytoplasm was subtracted from the background, and the ratio of centrosome/cytoplasm was used for plotting the graph. Signal intensities were compared using a two-sample unpaired t-test in GraphPad Prism.

### Chromosome condensation kinetics

The kinetics was done as described in (Maddox *et al*., 2006). Worms expressing GFP-H2B were recorded with z-stacking at 1.5 μm step size and 9 sections. Maximum z-projection was done on Fiji, and a rectangular region of interest around the chromosomes was cropped and converted to an 8-bit scale [0:255] at the time points indicated on the graph (Figure 2D). First, the total number of pixels in the ROI, then a threshold of 35% was applied, and the pixel count after thresholding was noted. The condensation parameter (Y-axis on Figure 2D) is defined as the percentage pixel count of the pixel number below 35% threshold divided by the total number of pixels in the ROI. Pixels *below* the applied threshold are proportional to chromosome condensation. The quantification pipeline was automated using a custom-built macro on Fiji to scale and threshold the image and calculate pixel count using the histogram function. Finally, the log values were normalized to 100% and arranged according to time series using MATLAB_R2020b.

## Biochemical assays

### Purification of GST-tagged LMN-1, LMN-1^H^, and LMN-1^T^ fragments

GST-LMN-1 was amplified from cDNA and cloned in pETM30 protein expression vector (6xHis-GST-TEV-LMN-1) and the purification was done as described (Velez-Aguilera et al., 2020). Protein expression was induced with 1mM IPTG in 400 ml of BL21 DE3 pLysS strain at OD=0.6-0.8, by incubation at 25°C for 3 hours. Cells pellet was collected by centrifugation at 6000 rpm for 10 minutes, and the pellet was then suspended in urea lysis buffer (8 M urea, 100 mM NaCl, 10 Mm Tris–HCl, pH 8, 1 mM 2-mercaptoethanol). Sonication was done at 30% amplitude, four times for 30 s each. Supernatant was collected after centrifugation at 11000 rpm for 30 minutes at room temperature. Purification was done using Ni-NTA (Qiagen: 30230) beads using gravity column, using binding buffer (8M Urea, 50mM Tris–HCl pH 8.0, 500mM NaCl, 20mM imidazole and 1 mM 2-mercaptoethanol). Beads were washed using wash buffer (8M Urea, 50mM Tris–HCl pH 8.0, 500mM NaCl, 40mM imidazole and 1 mM 2-mercaptoethanol). Elution was done in 8M urea, 10 mM Tris–HCl pH 8.0, 500mM imidazole, followed by step-wise dialysis to remove urea (10 mM Tris–HCl pH 8.0, 100 mM NaCl, 1 mM 2-mercaptoethanol). Protein dialysis was performed at 4°C and the protein was aliquoted and stored at -80°C after the snap freeze.

Likewise, LMN-1^H^ and LMN-1^T^ fragments were cloned in pETM30 protein expression vector (6xHis-GST-TEV-LMN-1^H^ and 6xHis-GST-TEV-LMN-1^T^). Protein expression was induced with 1mM IPTG in 400 ml of BL21 DE3 pLys cells at OD=0.6-0.8, by incubation at 25°C for 3 h. Cells pellet was collected by centrifugation at 5000 rpm for 20 minutes at 4°C and resuspended in lysis buffer (0.5M NaCl, 5% glycerol, 50mM Tris/HCl (pH=8), and 1x protease inhibitor cocktail). Lysozyme (Sigma, 1.5mg/ml) and DNase was added before sonication at 30% amplitude (3 s ON, 5 s OFF, 4-5 minutes). 0.5% Triton-X was added and the supernatant was collected after centrifugation at 13,000 rpm for 15 minutes at 4°C. For protein purification, the supernatant was incubated with NiNTA Beads (Qiagen: 30230) and 20mM Imidazole and kept at end-on rotation at 4°C. The supernatant was loaded on gravity column and washed with Wash Buffer (0.5M NaCl, 5% Glycerol, 50mM Tris-HCl (pH=8), 1x protease inhibitor cocktail, 40mM Imidazole). Bound protein was eluted in elution buffer containing 500mM Imidazole (100mM NaCl, 5% Glycerol, 10mM Tris-HCl (pH=8), and 1x protease inhibitor cocktail). Overnight protein dialysis (10 mM Tris–HCl pH 8.0, 100 mM NaCl, 1 mM 2-mercaptoethanol) was performed at 4°C and the protein was aliquoted and stored at -8°C after snap freeze.

### GFP-trap experiments

2 mg equivalent embryonic lysate from worms expressing GFP-B55^SUR-6^ was incubated with either 250 ng GST-LMN-1, or GST-LMN-1^H^, or GST-LMN-1^T^ with 0.2 mM working concentration of ATP at room temperature for 15 min. This was done to allow phosphorylation or modification, if any, of exogenously produced bacterial protein with embryonic lysate. An aliquot was collected as input fraction after this brief incubation, and the remaining sample was incubated with 25 ul GFP-Trap beads (Chromotek: ACT-CM-GFA0050). Before incubation, GFP-Trap beads were washed thrice in 500 ul ice-cold dilution buffer (lysis buffer without Glycerol and NP40) with DTT and PI (Merck: 539134 Merck) by spinning at 1000xg for 2 min. The beads were incubated with embryo lysate with GST-LMN-1, GST-LMN-1^H^, or GST-LMN-1^T^ and kept for end on rotation for 2-3 h at 4°C. The samples were spun at 2500xg for 2 min and the supernatant un-bound fraction was collected. 3 washes were done with dilution buffer followed by addition of 80 ul 2x SDS dye. The samples were boiled at 95°C for 10 min. The samples were spun at 2500xg for 5 minutes to collect the supernatant fraction, followed by immunoblotting.

### Immunoblotting

For immunoblotting, the samples were loaded on 8% SDS gel and transferred onto the nitrocellulose membrane (Biorad: 1620115) for immunoblot analysis. Blocking was done in BSA, and incubation with primary antibodies was done overnight at 40C. Anti-GST (Sigma: G7781) at 1:1000 dilution prepared in 5% BSA was used to detect LMN-1 protein. GFP-B55^SUR-6^ signal in the bead-bound fraction was detected using anti-FLAG (Sigma: F7425 Sigma) at 1:1000 prepared in 5% skimmed milk (Himedia: GRM1254). Since LET-92 protein sequence is 88.67% identical to human PPP2CA, PP2A C (CST: 2038) antibodies were used to detect LET-92, which detects a single band on the SDS-PAGE at the right size, and this band disappears in *let-92 (RNAi)* condition (data not shown).

## Acknowledgements

We thank Arshad Desai, Lionel Pintard, Federico Pelisch, Sudha Kumari, and Anthony Hyman, and Caenorhabditis Genetics Center (CGC) for sharing reagents and worm strains. We are grateful to Arshad Desai, Lionel Pintard, Griselda Velez-Aguilera, Batool Ossareh-Nazari, Martin Lowe, Andrew Goryachev, K Subramaniam, and the members of Kotak Laboratory for their critical comments on the manuscript. We thank DST-FIST, UGC Centre for the Advanced Study, Department of Biotechnology-Indian Institute of Science (DBT-IISc) Partnership Program, and IISc for the infrastructure support. This work is supported by the DBT grant (BT/PR36084/BRB/10/1857/2020) to S. Kotak, and by grants from the DBT/Wellcome Trust India Alliance Fellowship (IA/I/15/2/502077) to S. Kotak.

---

**Supplementary Figure 1.**
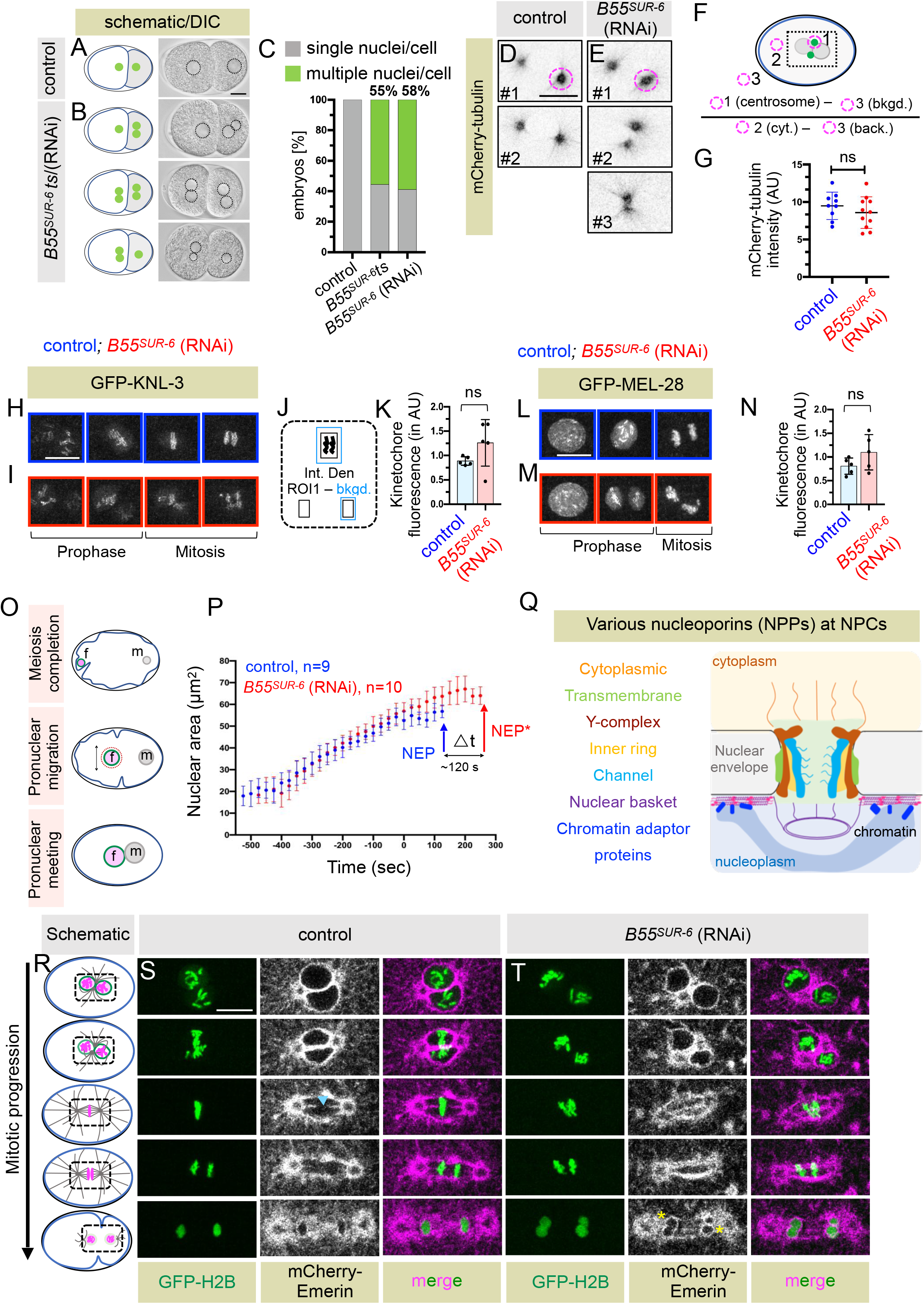
B55^SUR-6^ depletion or inactivation results in the formation of multiple nuclei in the two-cell stage embryo. (A-C) Schematic representation and the corresponding images from the time-lapse Differential Interference Contrast (DIC) microscopy of two-cell stage wild-type control (A) or *B55^SUR-6^ts/B55^SUR-6^ (RNAi)* (B) embryos. The nuclei have been marked by black dashed circles on the DIC images. The scale Bar is 10 μm unless specified in this and other Figure panels. The bar graph on the right (C) depicts the [%] of embryos that show either a single nucleus in each cell of the 2-cell stage embryos (as in 100% of the wild-type control embryo) or the presence of more than one nucleus per cell as depicted in B in *B55^SUR-6^ts* or *B55^SUR-6^ (RNAi)* embryos. [n= 15/27, 55.5% upon short and long-term upshift in *B55^SUR-6^ts* and n=10/17, 58.8% in *B55^SUR-6^ (RNAi)* embryos). (D, E) Images from the time-lapse confocal microscopy from different z-sections of embryos expressing mCherry-tubulin in control (D) and *B55^SUR-6^ (RNAi)* (E) at one frame before NEP, which was monitored by assessing the entry of mCherry-tubulin signal in the paternal nuclei. (F, G) Schematic representation of the quantification method (F) and the fluorescence intensity (G) of the mCherry-tubulin signal at the centrosome in control and *B55^SUR-6^ (RNAi)* embryos at one frame before NEP. bkgd., and cyt. represent background, and cytoplasmic intensity, respectively. Error bars SD; ns=0.315 as determined by two-tailed unpaired Student’s t-test. (H-K) Single nuclear insets from the z-sections of time-lapse confocal microscopy of embryos expressing GFP-KNL-3 in control (H) and *B55^SUR-6^ (RNAi)* embryos (I) at different stages of mitotic progression as indicated. Schematic representation of the quantification method (J) and the fluorescence intensity of GFP-KNL-3 at the kinetochore in control and *B55^SUR-6^ (RNAi)* embryos (K). In brief, the GFP intensity at the kinetochore has been quantified by subtracting background ROI from ROI placed at the kinetochore at the metaphase stage in maximum-z projections. Error bars SD; ns=0.125, as determined by two-tailed unpaired Student’s t-test. (L-N) Single nuclear insets from the z-sections of time-lapse confocal microscopy of embryos expressing GFP-MEL-28 in control and *B55^SUR-6^ (RNAi)* embryos at different stages of mitotic progression as indicated. The GFP-MEL-28 fluorescence intensity at the kinetochore has been quantified (N) similarly to the above Figure panel (K). Error bars SD; ns=0.150 as determined by two-tailed unpaired Student’s t-test. (O, P) Schematics illustrate different stages of cell cycle progression (O) and the quantification of the expansion of female pronuclear size in control (blue circles) and *B55^SUR-6^ (RNAi)* (red circles) embryos (P). The pronuclear meeting is time ‘0’, which remains unchanged in control and B55^SUR-6^*-*depleted embryos. Note that the NEP in *B55^SUR-6^ (RNAi)* embryos is delayed for ∼120s compared to control embryos with respect to the pronuclear meeting. (Q) Schematic representation of the nucleoporin complex (NPC) and the position of nucleoporins (color-coded) in NPC. Lamin is shown in pink, and the chromatin and chromatin adaptor-proteins are in blue. (R-T) Schematics illustrate different stages of the mitotic progression (R) with the nuclear insets in the schematic corresponding to the images from time-lapse confocal microscopy of embryos expressing mCherry-Emerin (grey); GFP-H2B (green) in control (S) and in *B55^SUR-6^ (RNAi)* (T) embryos. Note the occurrence of pronuclear scission monitored by the clearing of Emerin at adjoining regions of the male and female pronucleus in control embryos, indicated by blue arrowheads. In contrast, the mCherry-Emerin signal does not disappear in *B55^SUR-6^ (RNAi)* embryos; also note the formation of twin nuclei (shown by the yellow asterisk) at the two-cell stage in a significant number (∼38%) of *B55^SUR-6^* (RNAi) embryos in contrast to 0% of control embryos.

**Supplementary Figure 2.**
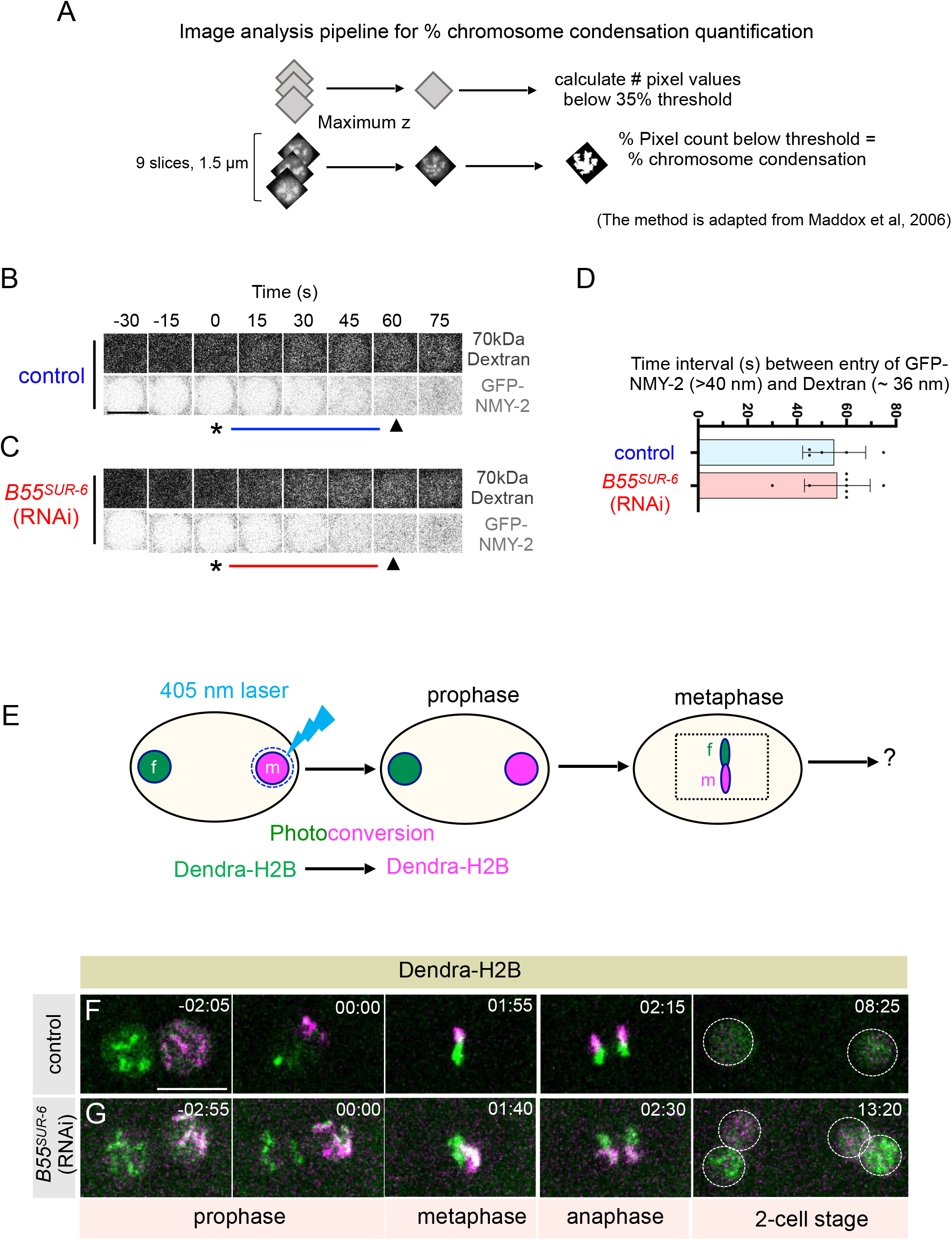
Paternal genomes fail to merge in B55^SUR-6^-depleted embryos. (A) Outline of the imaging pipeline used to analyze chromosome condensation (adapted from Maddox et al., 2006). Maximum z-projections of the nucleus are thresholded in 0:255 8-bit format, and the pixels below the threshold represent the percentage of chromosome condensation. 35% threshold has been used in the plot shown in Figure 2D. Also, see Methods for the detailed procedure. (B, C) Representative nuclear insets of the control (B) and *B55^SUR-6^ (RNAi)* (C) in GFP-NMY-2 expressing embryos from time-lapse confocal recordings of worms that are also injected with 70kDa Tetramethylrhodamine (TMR)-conjugated Dextran. The frame at which Dextran and GFP-NMY-2 enter the nucleus is indicated with black asterisk and black arrowhead in control and *B55^SUR-6^ (RNAi)* embryos. Time ‘0’ in (s) is relative to the entry of TMR-Dextran; The scale Bar is 10 μm in this and other Figure panels. (D) Bar graph represents the average time interval between the entry of TMR-Dextran and GFP-NMY-2 in control (blue, n=5) and *B55^SUR-6^* (RNAi) embryos (red, n=8). Error bars SD; ns=0.870 as determined by two-tailed unpaired Student’s t-test. (E-G) Schematics illustrate the photoconversion experiment performed in embryos expressing photoconvertible Denda-H2B. The male pronucleus was converted to magenta by exposure to UV light. The pattern of chromosome segregation of the male (magenta) and female (green) genomes was assessed by conducting time-lapse confocal microscopy. Maximum z-projections of nuclear insets from embryos expressing photoconvertible Denda-H2B in control (F) and *B55^SUR-6^* (RNAi) (G) embryos. Note the merging of magenta (male) and green (female) fluorescence signals at the two-cell stage in control in contrast to *B55^SUR-6^ (RNAi)* embryos. The nuclear insets are shown at prophase, metaphase, anaphase, and two-cell stage. Time ‘00.00’ is NEP, the expulsion of nucleoplasmic H2B signal from the nucleus. 6 embryos were recorded for control and *B55^SUR-6^* (RNAi) condition and the representative images are shown here.

**Supplementary Figure 3.**
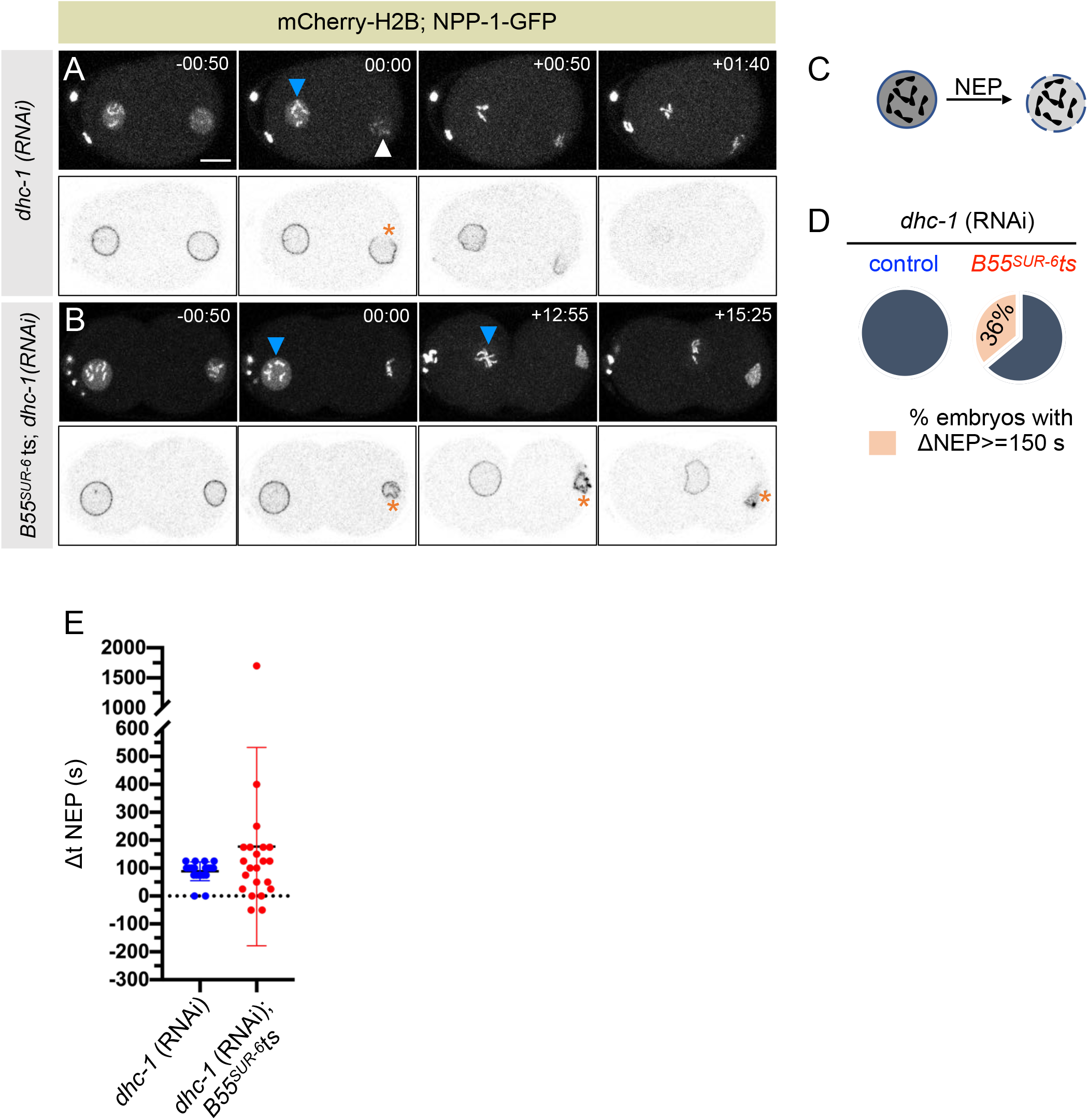
Dynein-dependent forces cooperate with B55^SUR-6^ for efficient NE dissolution. (A, B) Selected images from the time-lapse confocal microscopy of control (A) or *B55^SUR-6^ts* (B) embryos that are expressing NPP-1-GFP; mCherry-H2B and are depleted for dynein using *dhc-1 (RNAi)*. Time ‘00.00’ is the frame at which the male nucleus undergoes NEP. The nuclear wrinkling and H2B loss of the female pronucleus are excessively delayed in embryos co-depleted of DHC-1 and B55^SUR-6^, in contrast to control, DHC-1-depleted embryos alone. Scale bar 10 μm. (C) NEP was measured as rapid expulsion of the H2B signal shown in the schematic and used for quantification and analysis represented in (D, E). (D) Pie-chart depicting the pooled percentage of embryos showing asynchronous NEP (ΔNEP) in *dhc-1* (RNAi) control (blue circles) or *dhc-1* (RNAi) in *B55^SUR-6^ts* embryos at restrictive temperature (red circles). Note that ∼36% of the embryos in DHC-1 and B55^SUR-6^ co-depletion scenario show delayed asynchrony (DNEP>= 150 s, n=8/22). Error bars SD. (E) Quantification of the interval (in s) between permeabilization of the male and female pronuclei (referred to as ΔNEP) in *dhc-1 (RNAi)* control (blue circles) or *B55^SUR-6^ts* + *dhc-1 (RNAi)* (red circles) embryos. Note an exaggerated ΔNEP in *B55^SUR-6^(RNAi)/B55^SUR-6^ts* + *dhc-1 (RNAi)* embryos. Error bars SD.

**Supplementary Figure 4.**
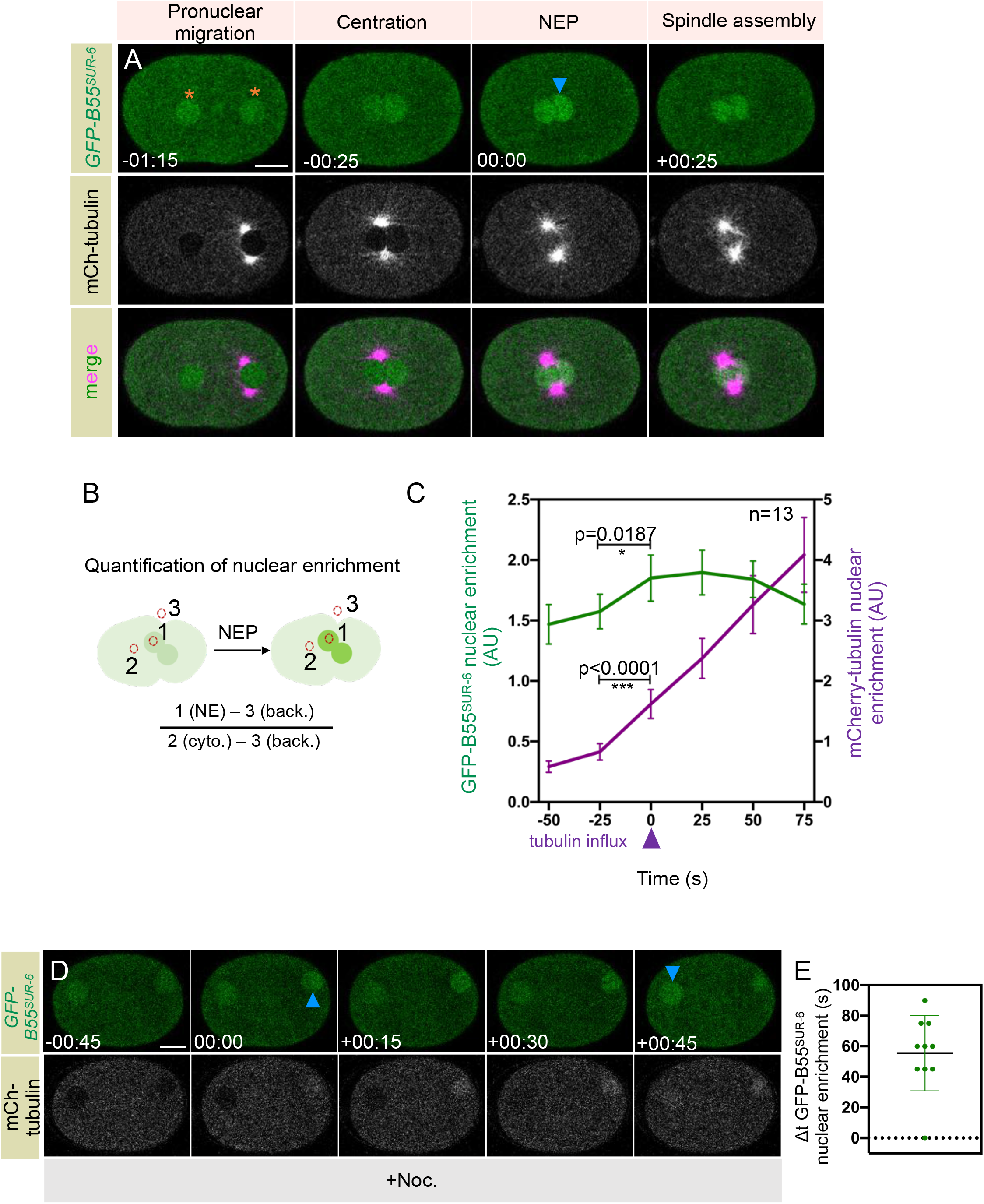
B55^SUR-6^ is rapidly imported into the nucleus at NEBD onset. (A) Selected images from the time-lapse confocal microscopy of embryos expressing endogenously-tagged B55^SUR-6^ with GFP and FLAG tag (referred to as GFP-B55^SUR-6^; in green) and mCherry-tubulin (in grey). Grayscale and the merged images show the acute entry of the GFP-B55^SUR-6^ signal, which coincides with the NEP (00:00) measured by the entry of the cytoplasmic tubulin signal in the nucleus. Orange asterisks depict the basel-level of nuclear enrichment of GFP-B55^SUR-6^ before NEP. The scale bar is 10 μm unless specified in this and other Figure panels. (B, C) Schematic representation of the quantification method (B) used to calculate nuclear enrichment of GFP-B55^SUR-6^ (green) and mCherry-tubulin (magenta) signal over cytoplasm. The average fluorescence intensity of nuclear enrichment of GFP-B55^SUR-6^ (green) and mCherry-tubulin (magenta) is plotted in (C) over time. Time ‘0’ is defined by the nuclear entry of the free tubulin inside the male/female pronucleus. Error bars SD; *P=0.0187 for the nuclear fluorescence intensity at the NEP with respect to one frame (25 s) before the NEP onset as determined by two-tailed unpaired Student’s t-test. (D, E) Selected images from the time-lapse confocal microscopy of embryos expressing GFP-B55^SUR-6^ (in green) and mCherry-tubulin (in grey), which are in addition treated with nocodazole. Time ‘00:00’ is defined as male NEP when soluble tubulin enters the male pronucleus. Note an average asynchrony (55.5 s) between the entry of the GFP-B55^SUR-6^ signal between the male and female pronucleus, shown in the scatter plot in (E). Dt is the average time interval of GFP entry in the female (late) versus the male (early) pronucleus. Error bars SD.

**Supplementary Figure 5.**
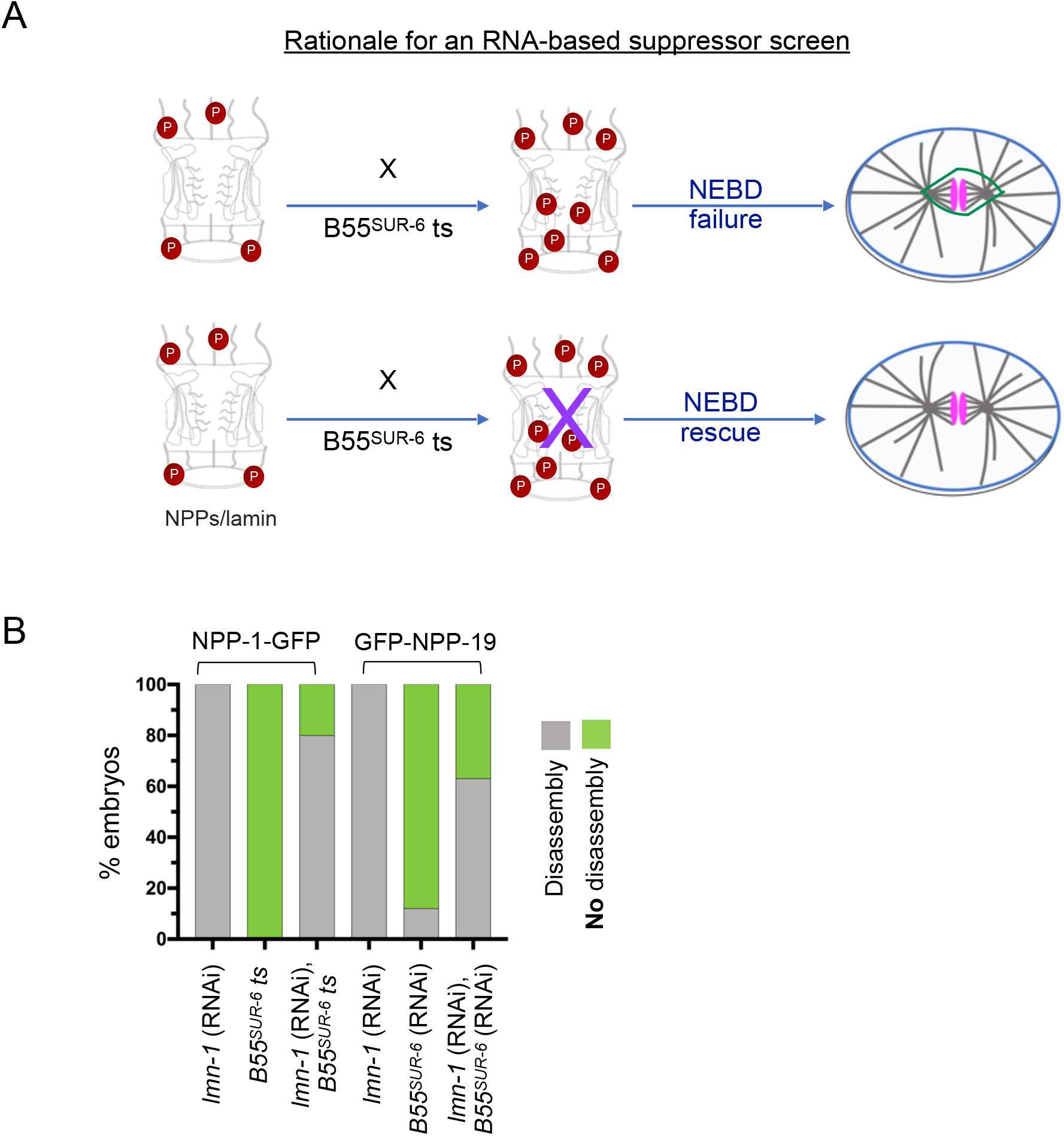
LMN-1 acts downstream of B55^SUR-6^ in regulation NEBD. (A) Schematic representation of the rationale for conducting a genetic suppressor screen. Co-depletion of B55^SUR-6^ with target protein such as NPP/(s) or the lamina, may result in the rescue of the nuclear envelope non-disassembly phenotype observed upon depletion of B55^SUR-6^. The working hypothesis is that hyper-phosphorylation of the target in the absence of B55^SUR-6^ would impair the timely nuclear envelope disassembly. Hence, the removal of the target could suppress the nuclear envelope disassembly defect. See Supplementary Table S1 for the nucleoporins used in the screen and the observations made upon double depletion of individual NPPs with B55^SUR-6^. (B) Bar graphs depicting rescue in the non-disassembly of nucleoporins observed upon B55^SUR-6^ loss. 80% of embryos that are depleted of LMN-1 in *B55^SUR-6^ ts* mutant background show dissolution of NPP-1-GFP signal [see Figure 5A-C], when compared to 0% *B55^SUR-6^ts* mutant embryos. Significant rescue (∼63%, n=10/17) was also seen in embryos expressing GFP-NPP-19 depleted of both LMN-1 and B55^SUR-6^ by RNAi (n=7, control, *lmn-1* (RNAi) embryos and n=17, *lmn-1*; *B55^SUR-6^* (RNAi) embryos).

