## Supplementary Table 1 for "PP2A-B55^SUR-6^ promotes nuclear envelope breakdown in *C. elegans* embryos"

| Supplementary Table 1 – List of RNAi used for feeding, and the outcome of the genetic suppressor screen | | |
|---|---|---|
| **List of RNAi** | **RNAi libraries [VL (Vidal Lab)* and AR (Ahringer Lab)*]** | **Condition** |
| *B55*$^{SUR\text{-}6}$ | VL (Vidal) and AR (Ahringer) | 24-48 hours |
| *lmn-1* | VL | 36-48 hours |
| *dhc-1* | VL | 48-72 hours |
| **List of Nucleoporin (NPP) RNAi** | **RNAi library** | **NPP-1-GFP (or YFP-LMN-1) signal in NPPs and *B55*$^{SUR\text{-}6}$ double (RNAi) embryos** |
| Y-complex | | |
| mel-28 | AR | Involved in nuclear assembly and hence excluded from analysis |
| npp-2 | VL | small, punctate pronuclei; No recue; Phenotype is *B55*$^{SUR\text{-}6}$ like |
| npp-5 | AR | No recue; Phenotype is *B55*$^{SUR\text{-}6}$ like |
| npp-6 | AR (12 hours short RNAi) | Punctate nuclei, No rescue |
| npp-10 | AR | Pronuclei formation was impaired; hence excluded from analysis |
| npp-15 | AR | No rescue; Phenotype is *B55*$^{SUR\text{-}6}$ like |
| npp-18 | VL | No rescue; Phenotype is *B55*$^{SUR\text{-}6}$ like |
| npp-20 | VL and AR | Dead embryos; hence excluded from analysis |
| npp-23 | VL | No rescue; Phenotype is *B55*$^{SUR\text{-}6}$ like |
| Cytoplasmic | | |
| npp-9 | AR (24 hours short RNAi) | No rescue; severely deformed nuclei because of npp-9 loss |
| npp-14 | VL | No rescue |
| npp-17 | VL | No rescue |
| npp-26/gle-1 | VL | No rescue; Phenotype is *B55*$^{SUR\text{-}6}$ like |

| | | |
|---|---|---|
| npp-24 | VL | No rescue; Phenotype is *B55*$^{SUR-6}$ like |
| Transmembrane | | |
| npp-12 | This study; cloned in L4440 | No rescue; Phenotype is sur-6-like |
| npp-22 | This study; cloned in L4440 | No rescue; Phenotype is sur-6-like |
| npp-25 | This study; cloned in L4440 | No rescue; Phenotype is sur-6-like |
| Inner ring | | |
| npp-3 | This study; cloned in L4440 | No rescue |
| npp-8 | This study; cloned in L4440 | Pronuclei defects; hence excluded from analysis |
| npp-10 | AR | Mentioned above in the cyto- and nucleoplasmic category |
| npp-13 | not included in the screen; could not source it from either library | |
| npp-19 | AR | No rescue |
| Central channel | | |
| npp-1 | This study; cloned in L4440 | Done in YFP-LMN-1 expressing embryos; No rescue |
| npp-4 | VL and AR | No rescue |
| npp-11 | This study; cloned in L4440 | No rescue |
| Nuclear basket | | |
| npp-7 | AR | Punctate, small pronuclei; no rescue |
| npp-16 | VL | No rescue; Phenotype is *B55*$^{SUR-6}$ like |
| npp-21 | AR | No rescue; Phenotype is *B55*$^{SUR-6}$ like |

Footnotes – Text in **bold** specifies header information
Highlighted text in grey indicates the subgroup of NPP

*VL – RNAi Library from Vidal Lab (Rual et al., 2004)
*AR – RNAi Library from Ahringer Lab (source BioScience, Kamath et al., 2003)
