## Supplementary Table 2 for "PP2A-B55^SUR-6^ promotes nuclear envelope breakdown in *C. elegans* embryos"

| Supplementary Table 2 – List of strains used in the study | | | | | |
|---|---|---|---|---|---|
| Strain | Identifier | Genotype | Source | Reference | Additional Information |
| N2 | *C. elegans* N2 Bristol | | CGC | | |
| GFP-B55$^{SUR-6}$ | OD4579 | | | Bel Borja et al., 2020 | Arshad Desai |
| PLK-1-GFP | OD2425 | plk-1(lt17) [plk-1::sgfp]loxp)III | CGC | Martino et al., 2017 | |
| YFP-LMN-1; mCherry-H2B | OD139 | ltIs37 [pie-1p::mCherry::his-58 + unc-119(+)] IV. qaIs3502[pie-1::YFP::LMN-1 + unc-119(+)] | CGC | | |
| NPP-1-GFP; mCherry-H2B | OCF3 | jjIs1092 [(pNUT1) npp-1::GFP + unc-119(+)]. ltIs37 [pie-1p::mCherry::his-58 + unc-119(+)] IV | CGC | Golden et al., 2009 | |
| GFP-H2B; mCherry-Emerin | it868 | | | | KS Subramanium |
| GFP-H2B; mCherry-SP12 | it907 | | | | KS Subramanium |
| Dendra-2-his-58/66 | OCF69 | ocfSi1 [mex-5p::Dendra2::his-58/66::tbb-2 3'UTR + unc-119(+)] I | CGC | Bolková and Lanctôt, 2016 | |
| GFP-MEL-28 | BN426 | mel-28(bq5[gfp::mel-28]) III | CGC | Gómez-Saldivar and Fernandez et al., 2016 | |
| GFP-NPP-9 | JH2184 | axIs1595 [pie-1p::GFP::npp-9(orf)::npp-9 3'UTR + unc-119(+)] | CGC | Voronina and Seydoux, 2010 | |
| GFP-NMY-2 | LP162 | cp13[nmy-2::gfp + LoxP] I | CGC | Dickinson et al., 2013 | |

| | | | | |
|---|---|---|---|---|
| GFP-LEM-2; mCherry-H2B | OD83 | ltIs37 [pie-1p::mCherry::his-58 + unc-119(+)] IV. qaIs3507 [pie-1::GFP::lem-2 + unc-119(+)] | CGC | |
| GFP-NPP-19 | BN46 | bqIs7 [pie-1p::LAP::npp-19 + unc-119(+)] | CGC | |
| GFP-KNL-3 | OD1 | ltIs1[pIC22; pie-1 promoter::knl-3::GFP + unc-119(+)] | CGC | |
| GFP-B55$^{SUR-6}$; mCherry tubulin | | | This study | |
| *B55$^{sur-6}$ ts (temperature sensitive)* | EU1062 | | CGC | O'Rourke et al., 2011 |
| *B55$^{sur-6}$ ts*; NPP-1-GFP; mCherry H2B | | | This study | |
| GFP-AIR-1; mCherry-PAR-2 | SK003 | | | Kapoor and Kotak, 2019 |
| GFP-TBB-2; mCherry TBG-1 | SA250 | tjIs54 [pie-1p::GFP::tbb-2 + pie-1p::2xmCherry::tbg-1 + unc-119(+)]. tjIs57 [pie-1p::mCherry::his-48 + unc-119(+)] | | Toya et al., 2010 |
